# Structural basis for piRNA-targeting

**DOI:** 10.1101/2020.12.07.413112

**Authors:** Todd A. Anzelon, Saikat Chowdhury, Siobhan M. Hughes, Yao Xiao, Gabriel C. Lander, Ian J. MacRae

## Abstract

Piwi proteins use PIWI-interacting RNAs (piRNAs) to identify and silence the transposable elements (TEs) pervasively found in animal genomes. The Piwi targeting mechanism is proposed to be similar to targeting by Argonaute proteins, which employ microRNA (miRNA) guides to repress cellular mRNAs, but has not been characterized in detail. We present cryo-EM structures of a Piwi-piRNA complex with and without target RNAs and analysis of target recognition. Resembling Argonaute, Piwi identifies targets using the piRNA seed-region. However, Piwi creates a much weaker seed so that prolonged target association requires further piRNA-target pairing. Beyond the seed, Piwi creates wide central cleft wide for unencumbered piRNA-target pairing, enabling long-lived Piwi-piRNA-target interactions that are tolerant of mismatches. Piwi ensures targeting fidelity by blocking propagation of the piRNA-target duplex in the absence of faithful seed pairing, and by requiring extended piRNA-target pairing to reach an endonucleolytically active conformation. This mechanism allows Piwi to minimize off-targeting cellular mRNAs and adapt piRNA sequences to evolving genomic threats.

## Introduction

PIWI-interacting RNAs (piRNAs) are the largest class of small (21-35 nt.) regulatory RNAs in animals, with tens of thousands to millions of unique piRNA species expressed in an individual organism (Wang, Zhang et al. 2019). piRNAs have diverse functions in animal biology, the most deeply conserved of which is preventing mobilization of transposable elements (TEs) (Ozata, Gainetdinov et al. 2019). piRNAs are ubiquitous throughout animal germlines, as failure to prevent TE-induced genome instability leads to partial or complete sterility (Rubin, Kidwell et al. 1982, Lin and Spradling 1997, Vagin, Sigova et al. 2006, Aravin, Sachidanandam et al. 2007, Brennecke, Aravin et al. 2007, Houwing, Kamminga et al. 2007, Kuramochi-Miyagawa, Watanabe et al. 2008, Khurana, Wang et al. 2011, Reuter, Berninger et al. 2011). Similarly, piRNA-associated factors are pervasive features of pluripotent stem cells in animals capable of extensive regeneration (Reddien, Oviedo et al. 2005, Rinkevich, Rosner et al. 2010, Zhu, Pao et al. 2012, Juliano, Reich et al. 2014). A wide variety of additional piRNA functions have been described, including acting as agents for passing epigenetic information from mother to embryo in fruit flies (Brennecke, Malone et al. 2008), determining gender in silkworms (Kiuchi, Koga et al. 2014), neutralizing viruses in mosquitos (Miesen, Joosten et al. 2016), and establishing long-term memories in sea slugs (Rajasethupathy, Antonov et al. 2012). However, functions for the majority of piRNA molecules found in most animals, including humans, are not known.

On the molecular level, piRNAs function through association with PIWI proteins (Aravin, Gaidatzis et al. 2006, Girard, Sachidanandam et al. 2006, Grivna, Beyret et al. 2006, Lau, Seto et al. 2006, Vagin, Sigova et al. 2006). PIWIs use piRNAs as guides to identify complementary sequences in RNAs targeted for silencing via a variety of mechanisms. PIWI proteins within the nucleus can direct histone/DNA methylation enzymes to silence genetic loci expressing transcripts recognized as piRNA targets (Aravin, Sachidanandam et al. 2008, Kuramochi-Miyagawa, Watanabe et al. 2008, Sienski, Donertas et al. 2012, Le Thomas, Rogers et al. 2013), while in the cytoplasm PIWI proteins have been proposed to recruit deadenylases to degrade messenger RNAs (mRNA) targeted by piRNAs (Rouget, Papin et al. 2010, Gou, Dai et al. 2014). Most PIWI proteins also use an intrinsic endonuclease activity to cleave target RNAs (Gunawardane, Saito et al. 2007, Reuter, Berninger et al. 2011), which can subsequently be further fragmented to generate new piRNAs. Such cleavage events lead to piRNA amplification and expansion of the cellular piRNA pool to incorporate additional sequences within target RNA molecules (Brennecke, Aravin et al. 2007, Gunawardane, Saito et al. 2007, Han, Wang et al. 2015, Homolka, Pandey et al. 2015, Mohn, Handler et al. 2015, Gainetdinov, Colpan et al. 2018). Therefore, piRNA-targeting, the mechanism by which the PIWI-piRNA complex identifies targets, shapes the cellular piRNA repertoire and profoundly impacts gene silencing.

Currently, small RNA targeting mechanisms are best understood for mammalian Argonaute2 (Ago2). Ago2 uses microRNAs (miRNAs), a second major class of animal small RNAs, to identify mRNA targets to be repressed as a normal part of health and development (Bartel 2018). Ago2 facilitates target recognition by pre-ordering the guide (g) nucleotides g2-g7 (referred to as the canonical “seed” region) of the miRNA, into a helical conformation and thereby lowers the entropic cost of target binding (Filipowicz 2005, Parker, Roe et al. 2005, Tomari and Zamore 2005, Wang, Sheng et al. 2008, Parker, Parizotto et al. 2009, Nakanishi, Weinberg et al. 2012, Schirle and MacRae 2012, Wee, Flores-Jasso et al. 2012, Nakanishi, Ascano et al. 2013, Park, Phan et al. 2017, Park, Araya-Secchi et al. 2019). Ago2 further shapes miRNA-target interactions by creating a central cleft that includes two chambers with space for target pairing to miRNA seed and supplementary (g13-g17) nucleotides, and two narrow regions, which discourage pairing to the miRNA central (g9–g12) and 3’ tail (g18– g22) nucleotides (Schirle, Sheu-Gruttadauria et al. 2014, Sheu-Gruttadauria, Xiao et al. 2019).

piRNAs are proposed to use a miRNA-like targeting mechanism (Gou, Dai et al. 2014, Shen, Chen et al. 2018). Supporting this notion, seed complementarity is a recurrent feature of piRNA target transcripts identified *in vivo* (Goh, Falciatori et al. 2015, Zhang, Kang et al. 2015, Shen, Chen et al. 2018, Zhang, Tu et al. 2018, Halbach, Miesen et al. 2020). Additionally, PIWI and Argonaute (AGO) family proteins share a common set of domains and have extensive sequence similarities (Carmell, Xuan et al. 2002). On the other hand, crystal structures of *Drosophila* Piwi and silkworm Siwi have notable differences in the three-dimensional arrangement of domains compared to known AGO structures, indicating that piRNA-target interactions may be shaped in a distinct manner (Matsumoto, Nishimasu et al. 2016, Yamaguchi, Oe et al. 2020). It has not been possible to distinguish between these possibilities and directly measure piRNA-target interactions because a homogenous source of Piwi-piRNA complex suitable for biochemical and structural analysis has not been available.

Here, we describe a recombinant source of Piwi-piRNA complex, derived from the *piwi-a* gene of the freshwater sponge *Ephydatia fluviatilis*. The protein (EfPiwi) can be produced in a baculovirus expression system and loaded with a synthetic piRNA, resulting in a stable RNA-protein complex that can be purified to homogeneity. We determined cryo-electron microscopy (cryo-EM) structures of the EfPiwi-piRNA complex with and without target RNAs and performed biochemical analyses of target recognition. Like mammalian Ago2, EfPiwi initially engages targets through seed pairing. However, EfPiwi creates a much weaker seed than Ago2 such that further base pairing is required for high-affinity target interactions. EfPiwi enforces targeting fidelity by extensively probing the piRNA-target duplex in the seed region and employing a structural element, termed the “seed-gate”, to block duplex propagation in the absence of faithful seed pairing. Beyond the seed-gate, EfPiwi creates central cleft wide enough for unencumbered piRNA-target pairing, enabling EfPiwi-piRNA-target interactions that are long-lived and tolerant of mismatches. Target cleavage, however, requires an extended piRNA-target duplex of sufficient length and rigidity to drive EfPiwi into a fully open conformation and activate endonuclease activity. We propose the overall targeting mechanism allows Piwi-piRNA complexes to accumulate on and/or destroy target transcripts a manner that is sufficiently specific to avoid off-targeting cellular RNAs yet malleable enough to adapt to new and evolving threats to the genome.

### Recombinant Source of homogenous PIWI-piRNA complexes

PIWI proteins derived from natural sources co-purify with heterogenous mixtures of endogenous piRNAs (Aravin, Gaidatzis et al. 2006, Girard, Sachidanandam et al. 2006, Grivna, Beyret et al. 2006, Lau, Seto et al. 2006, Vagin, Sigova et al. 2006, Kawaoka, Hayashi et al. 2009), which complicate biochemical and structural analysis of targeting. Therefore, we first sought to establish a recombinant system for preparing homogenous PIWI-piRNA complexes. Expression levels of recombinant PIWI proteins derived from various animals (including mouse, silk moth, fruit fly, worm, zebra fish, flour beetle, and sponge) were screened in a baculovirus system to identify constructs suitable for biochemical and structural analysis. A working construct was obtained using *piwi-a* from the freshwater sponge *Ephydatia fluviatilis* (Funayama, Nakatsukasa et al. 2010) (**Fig. 1A**). *Ephydatia fluviatilis* Piwi-A (hereafter referred to as EfPiwi) is highly enriched in totipotent archeocytes, which have the potential to generate both germ-line and somatic cells in the sponge (Alie, Hayashi et al. 2015). EfPiwi belongs to a deeply rooted branch of the PIWI family tree that includes *Drosophila* Ago3 and Piwi-like proteins MILI and HILI (Alie, Hayashi et al. 2015, Wynant, Santos et al. 2017, Jehn, Gebert et al. 2018), and is distinct from the insect-specific offshoot containing *Drosophila* Piwi (DmPiwi) and Siwi (**Fig. 1B**). The best expressing EfPiwi construct has a deletion of the first 219 amino acids, removing an N-terminal region that is divergent among PIWI members and predicted to be intrinsically disordered (Jones and Cozzetto 2015). Recombinant EfPiwi protein can be loaded with a chemically defined single-stranded exogenous piRNA, and the resulting piRISC complex can be purified using an immobilized complementary target oligonucleotide (Flores-Jasso, Salomon et al. 2013) to produce a homogenous sample (**Fig. 1C**). This system enables biochemical and structural analyses to an extent previously not possible for a PIWI-piRNA complex.

**Figure 1.**
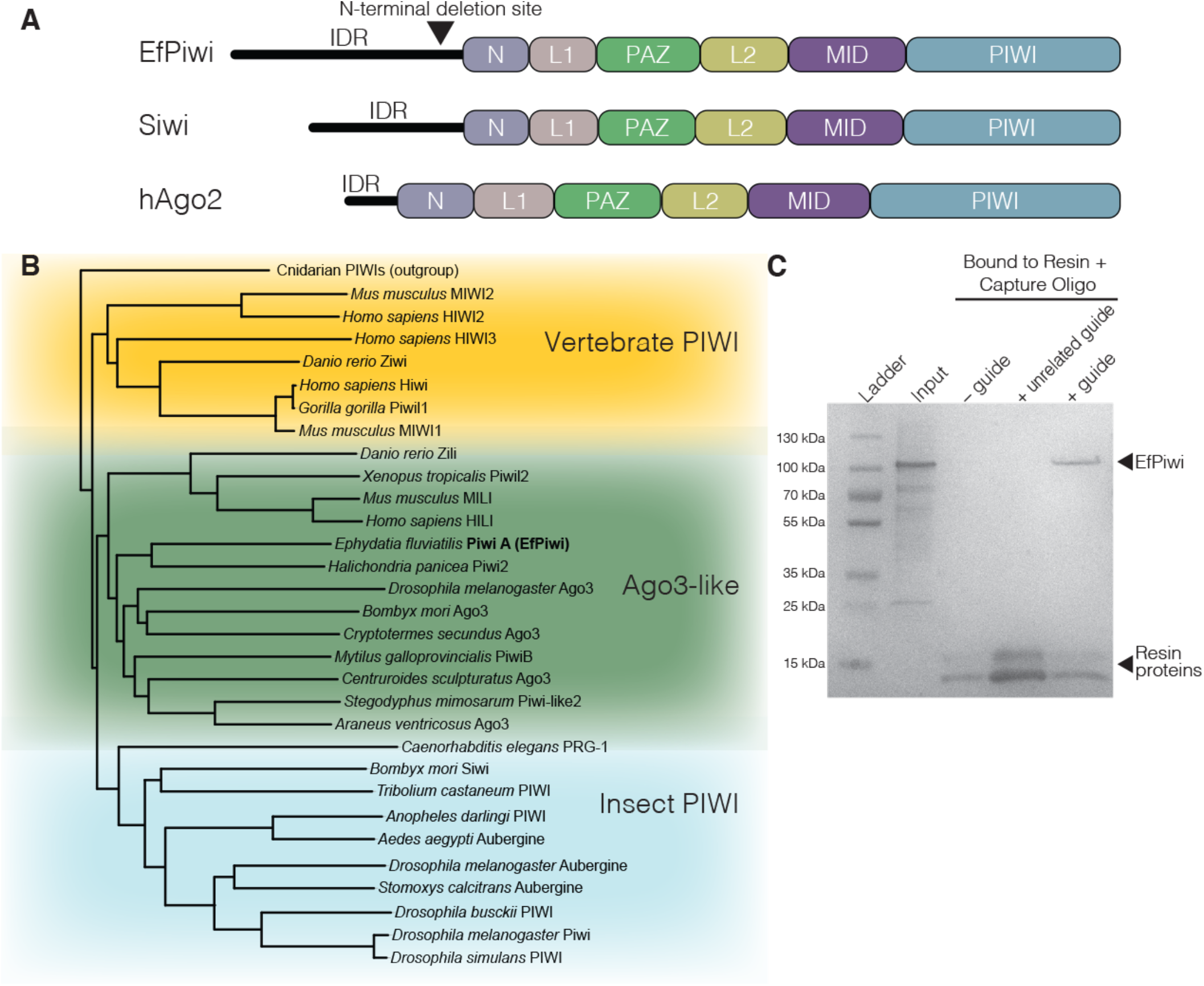
Recombinant Source of an Active PIWI Protein. **A**. Schematic of EfPiwi primary sequence with major domains labeled. Siwi and hAgo2 included for comparison. **B**. Phylogenetic tree of PIWI proteins shows EfPiwi belongs to the *Drosophila* AGO3-like branch. **C**. Coomassie-stained SDS-PAGE showing the EfPiwi-piRNA complex is specifically bound immobilized complementary target oligonucleotide (protein was eluted from resin by incubation boiling with SDS).

### Structure of the EfPiwi-piRNA complex reveals common features of the PIWI-clade

We used cryo-EM single-particle analysis to determine the structure of EfPiwi bound to a 25 nucleotide (nt) piRNA to an overall resolution of ∼3.8 Å (**Fig. 2A, Fig. S2**). The MID, PIWI, N, and PAZ domains known to be present in AGO and PIWI proteins were all observed. The reconstruction was of sufficient quality to build a full atomic model of the protein elements except for a few loops and the N domain, likely due to flexibility of these regions. Notably, the piRNA was mostly flexible with poorly defined density, with the exception of the 5’ and 3’ ends (guide nucleotides g1–g4 and g23–g25), which were ordered well enough to be modeled (**Fig. 2A and B, Fig. S2**). Weak density for a metal ion coordinated to the piRNA 5’ phosphate, as in crystal structures of Siwi and *Drosophila* Piwi (Matsumoto, Nishimasu et al. 2016, Yamaguchi, Oe et al. 2020), was also observed.

**Figure 2.**
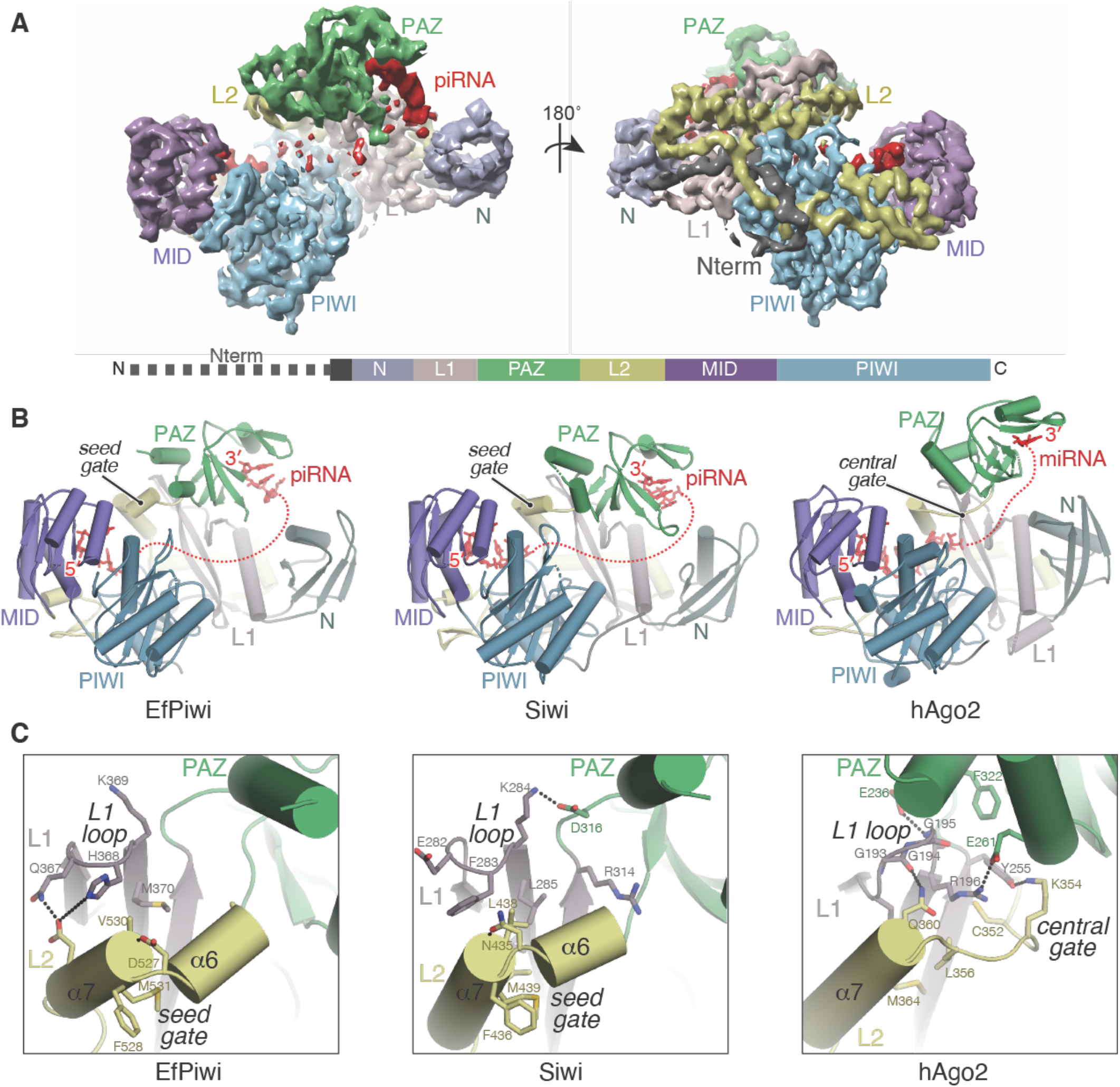
Cryo-EM structure of EfPiwi. **A**. Segmented cryo-EM reconstruction with major domains colored distinctly. Linear domain schematic is shown (lower panel). **B**. Cartoon representations of EfPiwi (left), Siwi (PDB 5GUH, middle), and hAgo2 (PDB 4OLA, right) shown in the same orientation with domains and major features indicated. Disordered regions of bound guide RNAs indicated by dashed red lines. **C**. Close-up view of the L1-PAZ-L2 interface of EfPiwi, Siwi, and Ago2 with structural interface residues shown.

The three-dimensional arrangement of domains in EfPiwi is similar to that of Siwi and *Drosophila* Piwi, although distinct from AGO-clade structures, such as human Ago2 (hAgo2) (**Fig. 2B**). Differences at the three-way interface of the L1, L2 and PAZ domains reposition residues that compose the Ago2 central gate, which restricts guide-target interactions in the miRNA central region (Schirle, Sheu-Gruttadauria et al. 2014, Sheu-Gruttadauria, Xiao et al. 2019), to a location adjacent to the piRNA seed region. Thus, EfPiwi contains an expanded ‘seed-gate’ structure, which reshapes the space near the piRNA seed region (**Fig. 2C**), and a widened central cleft compared to hAgo2. Siwi has similar features and the residues that compose the L1-L2-PAZ interface are conserved within the PIWI and AGO clades. These findings support the hypothesis that PIWI-clade proteins have common three-dimensional architectural features, which may shape target-recognition properties in a manner distinct from other eukaryotic Argonaute family members (Matsumoto, Nishimasu et al. 2016, Yamaguchi, Oe et al. 2020).

**Figure S2.**
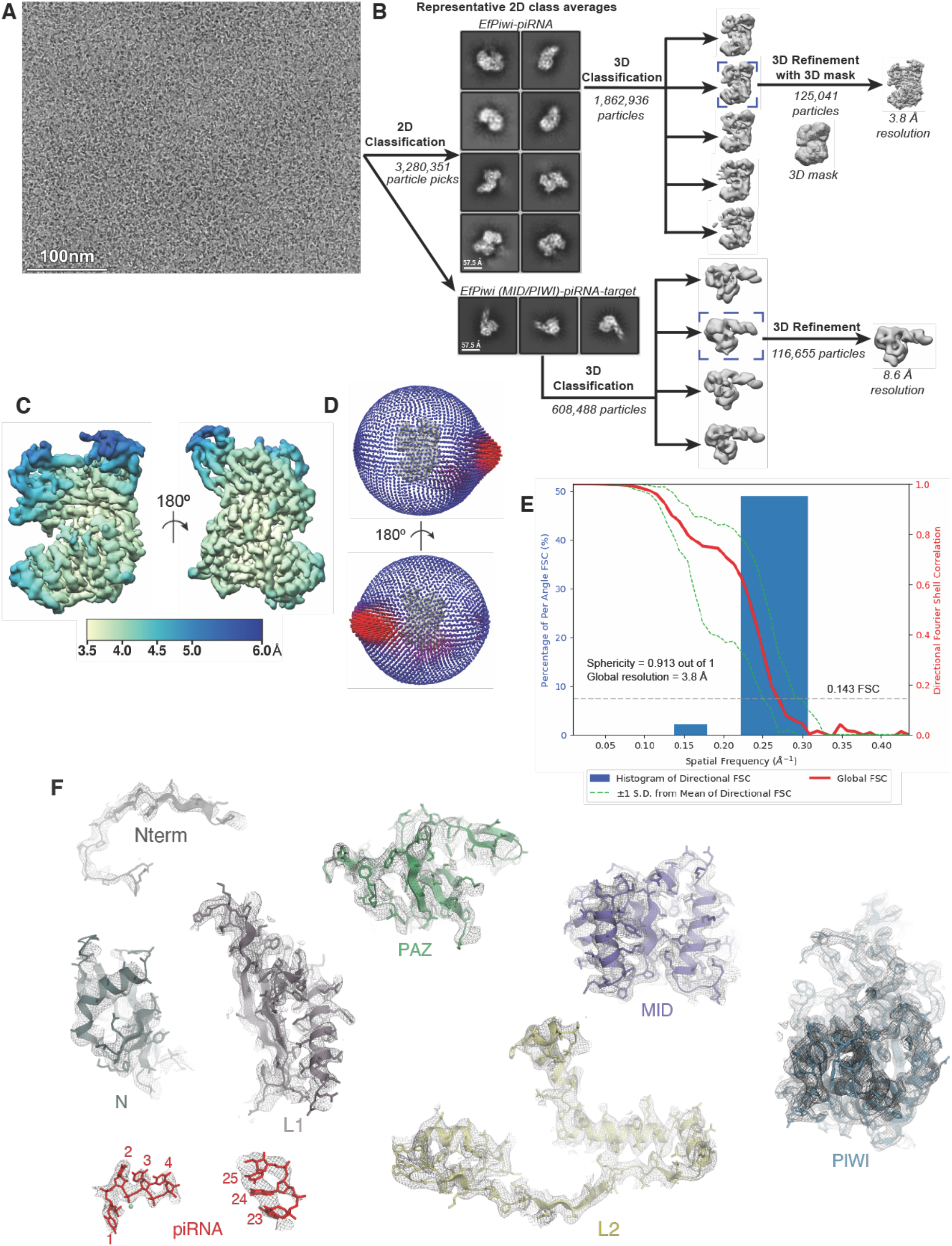
Imaging and processing of the EfPiwi-piRNA complex (and EfPiwi-piRNA-long target complex). **A**. Representative cryo-EM micrograph. Input sample contained EfPiwi-piRNA and a long target RNA (complementary to piRNA nucleotides g2–g25). **B**. Cryo-EM data processing workflow. The data set contained two populations of well resolved particles, one for the binary EfPiwi-piRNA complex and another for the ternary EfPiwi-piRNA-long target complex. Particles isolated from micrographs were sorted by reference-free 2D classification. Only particles containing high-resolution features for the intact complex were selected for downstream processing. 3D classification was used to further remove low-resolution or damaged particles, and the remaining particles were refined to obtain a 3.8 Å reconstruction for the EfPiwi-guide complex, and 8.6 Å for the ternary EfPiwi-piRNA-long target complex. **C**. The final 3D map for the EfPiwi-piRNA complex colored by local resolution values, where the majority of the map was resolved between 3.5 Å and 4 Å with the flexible PAZ and N domains having lower resolution. **D**. Angular distribution plot showing the Euler angle distribution of the EfPiwi-piRNA particles in the final reconstruction. The position of each cylinder corresponds to the 3D angular assignments and their height and color (blue to red) corresponds to the number of particles in that angular orientation. **E**. Directional Fourier Shell Correlation (FSC) plot representing 3D resolution anisotropy in the reconstructed map, with the red line showing the global FSC, green dashed lines correspond to ±1 standard deviation from mean of directional resolutions, and the blue histograms correspond to percentage of directional resolution over the 3D FSC. **F**. EM density quality of EfPiwi-piRNA complex. Individual domains of EfPiwi fit into the EM density, EM density shown in mesh; molecular models (colored as in Fig. 2) shown in cartoon representation with side chains shown as sticks; piRNA shown in stick representation.

**Table S1.**
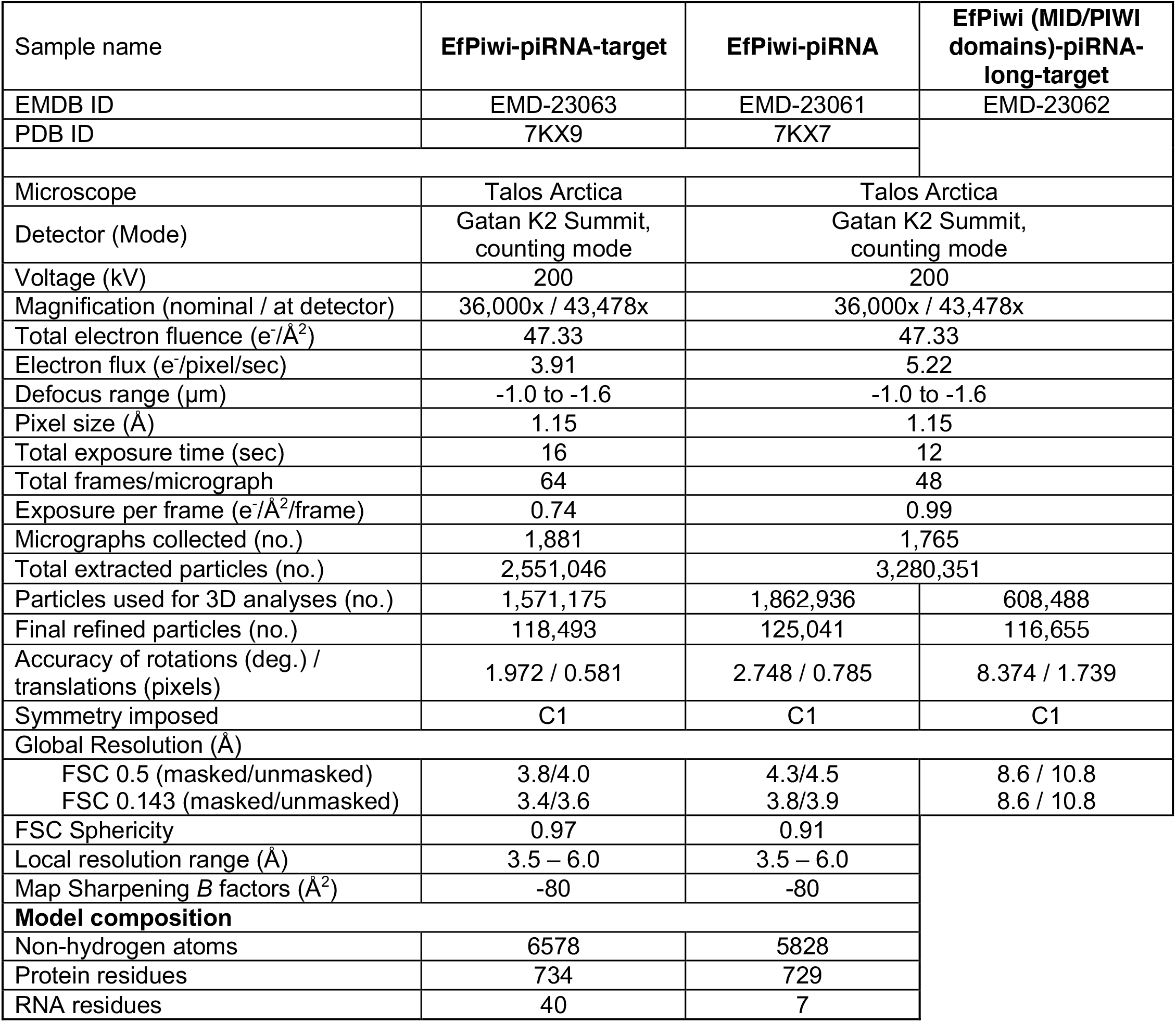
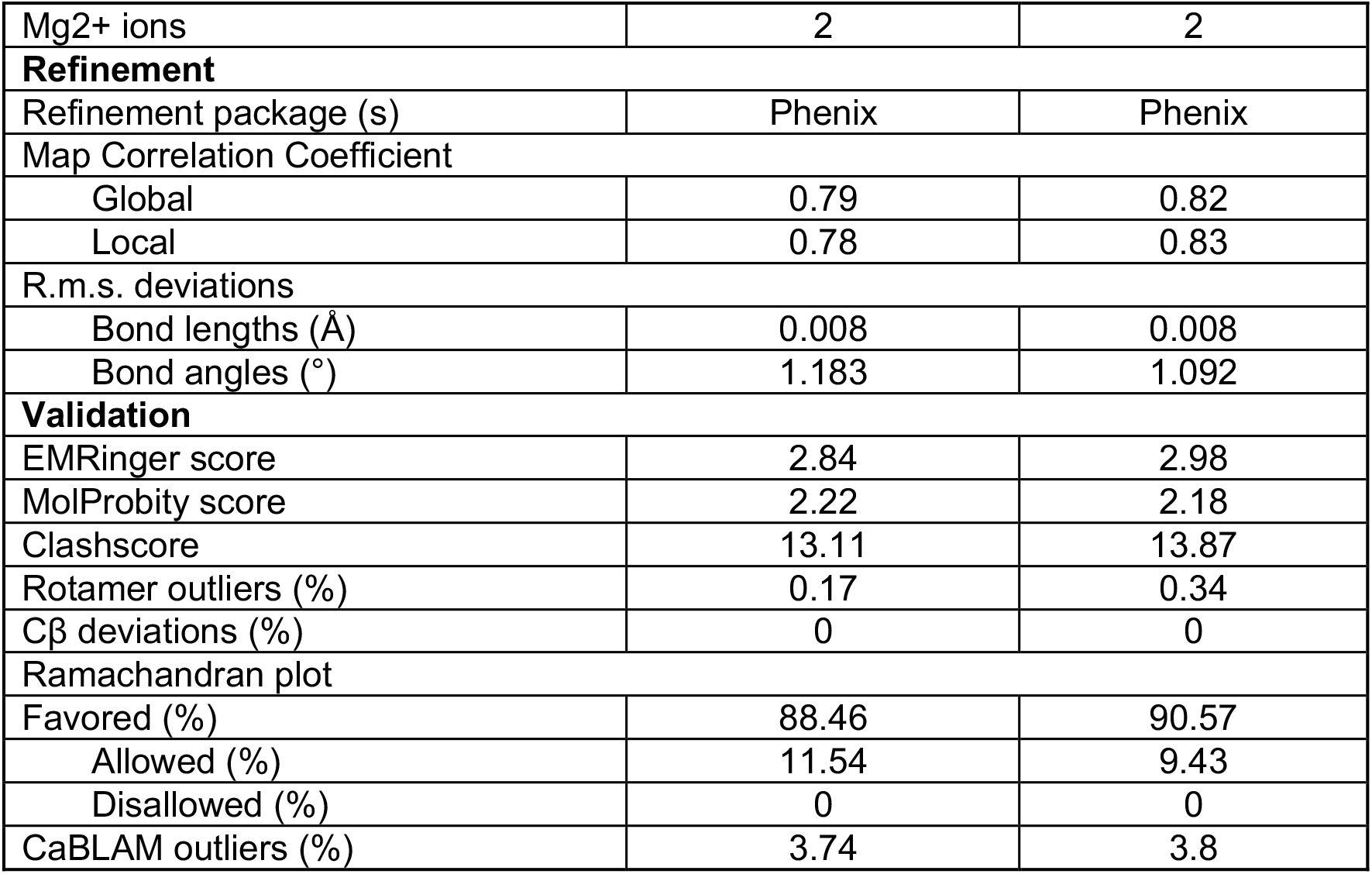
Cryo-EM data collection, refinement, and validation statistics.

| Sample name | EfPiwi-piRNA-target | EfPiwi-piRNA | EfPiwi (MID/PIWI domains)-piRNA-long-target |
| --- | --- | --- | --- |
| EMDB ID | EMD-23063 | EMD-23061 | EMD-23062 |
| PDB ID | 7KX9 | 7KX7 |  |
| Microscope | Talos Arctica | Talos Arctica |  |
| Detector (Mode) | Gatan K2 Summit, counting mode | Gatan K2 Summit, counting mode |  |
| Voltage (kV) | 200 | 200 |  |
| Magnification (nominal / at detector) | 36,000x / 43,478x | 36,000x / 43,478x |  |
| Total electron fluence ($e^-/\text{\AA}^2$) | 47.33 | 47.33 | |
| Electron flux ($e^-/\text{pixel}/\text{sec}$) | 3.91 | 5.22 | |
| Defocus range ($\mu\text{m}$) | -1.0 to -1.6 | -1.0 to -1.6 | |
| Pixel size ($\text{\AA}$) | 1.15 | 1.15 | |
| Total exposure time (sec) | 16 | 12 |  |
| Total frames/micrograph | 64 | 48 |  |
| Exposure per frame ($e^-/\text{\AA}^2/\text{frame}$) | 0.74 | 0.99 | |
| Micrographs collected (no.) | 1,881 | 1,765 |  |
| Total extracted particles (no.) | 2,551,046 | 3,280,351 |  |
| Particles used for 3D analyses (no.) | 1,571,175 | 1,862,936 | 608,488 |
| Final refined particles (no.) | 118,493 | 125,041 | 116,655 |
| Accuracy of rotations (deg.) / translations (pixels) | 1.972 / 0.581 | 2.748 / 0.785 | 8.374 / 1.739 |
| Symmetry imposed | C1 | C1 | C1 |
| Global Resolution ($\text{\AA}$) | | | |
| FSC 0.5 (masked/unmasked) | 3.8/4.0 | 4.3/4.5 | 8.6 / 10.8 |
| FSC 0.143 (masked/unmasked) | 3.4/3.6 | 3.8/3.9 | 8.6 / 10.8 |
| FSC Sphericity | 0.97 | 0.91 |  |
| Local resolution range ($\text{\AA}$) | 3.5 – 6.0 | 3.5 – 6.0 | |
| Map Sharpening $B$ factors ($\text{\AA}^2$) | -80 | -80 | |
| **Model composition** |  |  |  |
| Non-hydrogen atoms | 6578 | 5828 |  |
| Protein residues | 734 | 729 |  |
| RNA residues | 40 | 7 |  |

|  |  |  |
| --- | --- | --- |
| Mg<sup>2+</sup> ions | 2 | 2 |
| **Refinement** |  |  |
| Refinement package (s) | Phenix | Phenix |
| Map Correlation Coefficient |  |  |
| Global | 0.79 | 0.82 |
| Local | 0.78 | 0.83 |
| R.m.s. deviations |  |  |
| Bond lengths (Å) | 0.008 | 0.008 |
| Bond angles (°) | 1.183 | 1.092 |
| **Validation** |  |  |
| EMRinger score | 2.84 | 2.98 |
| MolProbity score | 2.22 | 2.18 |
| Clashscore | 13.11 | 13.87 |
| Rotamer outliers (%) | 0.17 | 0.34 |
| Cβ deviations (%) | 0 | 0 |
| Ramachandran plot |  |  |
| Favored (%) | 88.46 | 90.57 |
| Allowed (%) | 11.54 | 9.43 |
| Disallowed (%) | 0 | 0 |
| CaBLAM outliers (%) | 3.74 | 3.8 |

### Piwi creates a weak seed region

AGO proteins examined to date create a seed region that serves as the primary determinant of target binding affinity for miRNAs and small interfering RNAs (siRNAs) (Lai and Posakony 1998, Lai 2002, Haley and Zamore 2004, Brennecke, Stark et al. 2005, Krek, Grun et al. 2005, Lim, Lau et al. 2005, Wee, Flores-Jasso et al. 2012). AGOs form the seed by pre-organizing guide nucleotides g2–g7 in a helical conformation. We noticed, however, that EfPiwi pre-organizes only half (g2–g4) of the seed (**Fig. 3A**) and, even upon target binding (see below), makes half as many backbone contacts to the guide seed region as hAgo2 (**Fig. S3A**). We therefore hypothesized that target-binding properties of the piRNA seed may be distinct from those of miRNAs and siRNAs.

**Figure 3.**
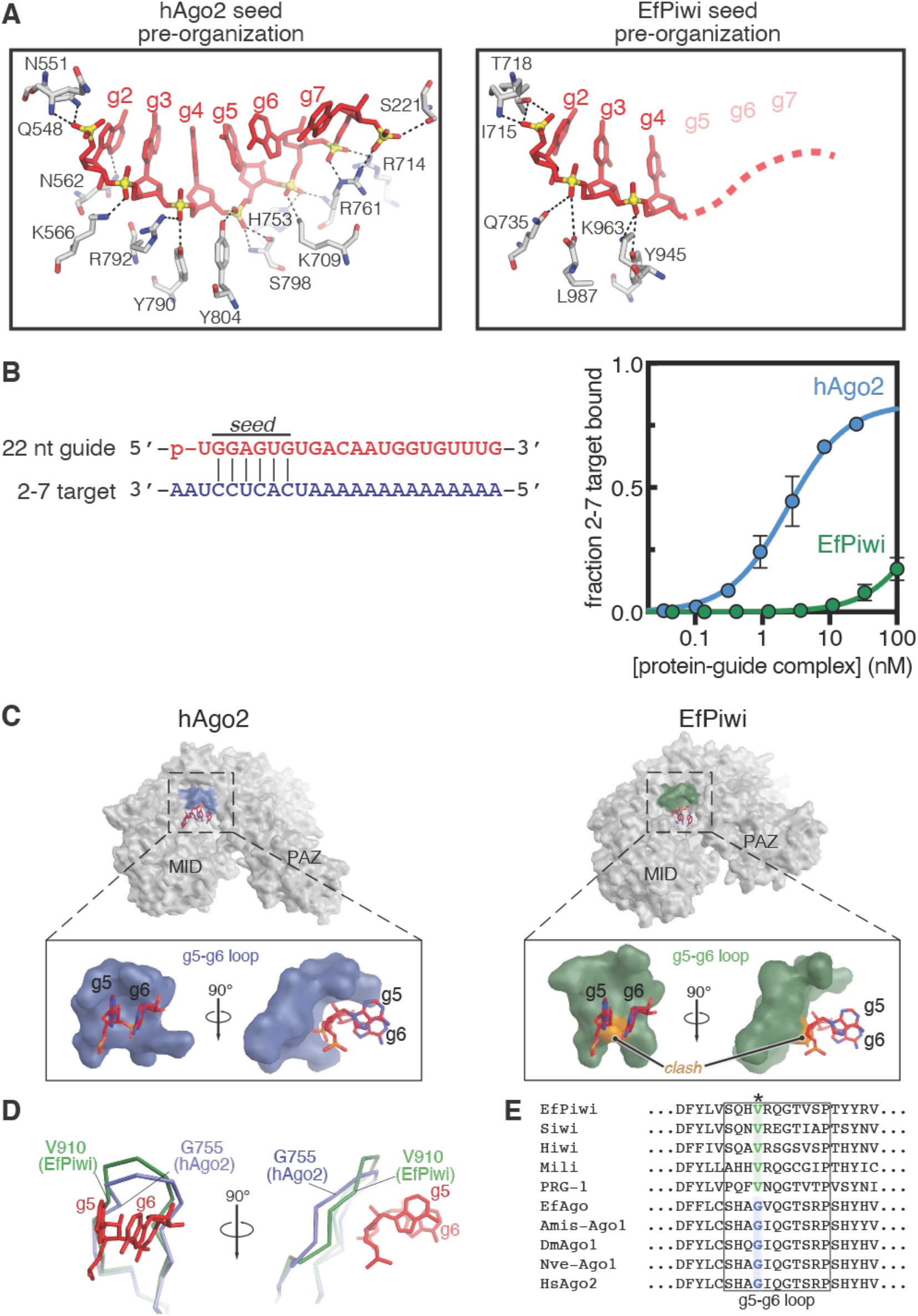
Piwi creates a weak canonical seed. **A**. Close up of miRNA (within hAgo2, left) and piRNA (within EfPiwi, right) seed regions prior to target binding. Amino acid residues contacting the RNAs are shown. Dashed red line indicates unstructured piRNA nucleotides. **B**. Schematic of seed-matched target RNA paired to guide (left) and relative fraction seed-matched target bound as a function of hAgo2-guide or EfPiwi-guide concentration. **C**. Surface representation highlighting the loop that cradles g5-g6 in hAgo2 (left), and the equivalent loop in EfPiwi (right). g5-g6 nucleotides from the hAgo2 structure are shown superimposed on EfPiwi. Atoms in EfPiwi that sterically clash with the modeled RNA are colored orange. **D**. Superposition of the g5-g6-cradeling loops shows a glycine residue in hAgo2 (G755) provides a kink necessary for cradling. **E**. Sequence alignment of AGO- and PIWI-clade proteins shows conservation of hAgo2 G755 within the AGO-clade, with PIWI-clade proteins possessing a valine at the equivalent position.

We compared the affinities of EfPiwi and hAgo2 for seed matched target RNAs. Both proteins were loaded with identical 22 nt guide RNAs and the resulting RNA-protein complexes were purified and incubated with a target with complementarity restricted to the canonical seed (g2-g7) (**Fig. 3B**). EfPiwi bound the seed-matched target with >150-fold lower affinity than Ago2 (*K*_*D*_ values of 399 ± 31.4 nM and 2.37 ± 0.08 nM for EfPiwi and Ago2, respectively) (**Fig. 3B**). The affinity of EfPiwi for the seed-matched target was not substantially changed by increasing guide length to 25 nt and including a 2′-O-methyl (2′-OCH_3_) group on the guide 3’ end in order to emulate a poriferan piRNA (Grimson, Srivastava et al. 2008) (**Fig. S3C and D**). Similarly, EfPiwi bound a target complementary to an ‘extended seed’ (g2–g8) (Bartel 2018) with 10-fold lower affinity, and a 10-fold faster release rate, than hAgo2 (**Fig. S3E**). We conclude EfPiwi creates a weaker seed than hAgo2.

Superimposing coordinates of seed region nucleotides from hAgo2 onto the EfPiwi structure results in a steric clash between guide nucleotides g5-g6 and a loop in EfPiwi (**Fig 3C**). hAgo2 avoids this clash through a glycine kink, which allows the loop to cradle g5-g6. EfPiwi contains a valine residue in place of the glycine and thus lacks the kink and ability to pre-organize the seed beyond g4 in a manner similar to hAgo2 (**Fig. 3D**). Glycine and valine residues are conserved at this position within AGO and PIWI families, respectively (**Fig. 3E**), and structures of Siwi and DmPiwi both show the equivalent loop does not cradle or pre-organize g5-g6 (Matsumoto, Nishimasu et al. 2016, Yamaguchi, Oe et al. 2020). We suggest weak canonical seed pairing may be a general feature of piRNAs.

**Figure S3.**
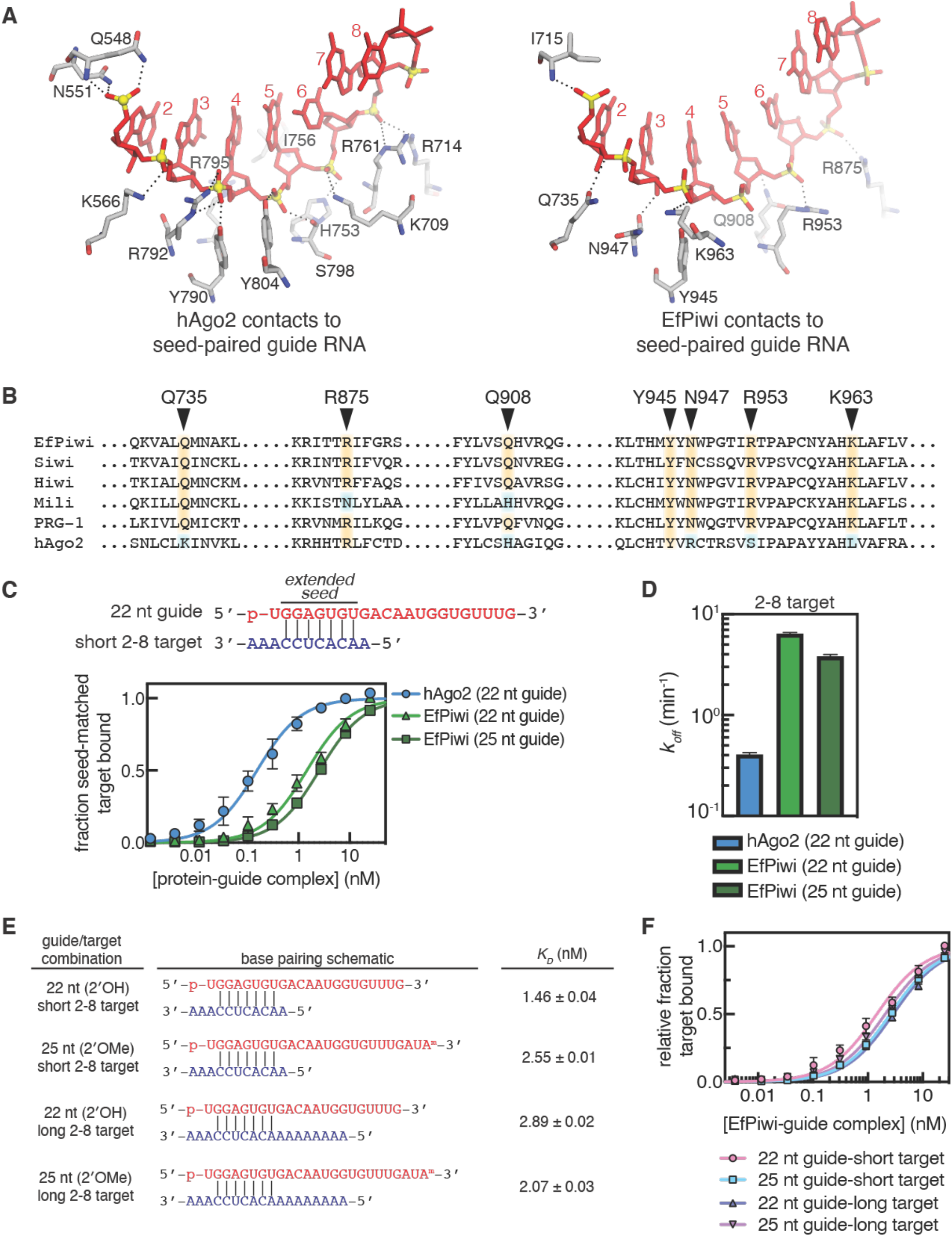
EfPiwi creates a weak seed region. **A**. Contacts between guide RNA seed region and hAgo2 (left) or EfPiwi (right) in target-bound structures. **B**. Sequence alignment showing conservation of seed-contacting amino acid residues within PIWI-clade. **C**. Schematic of short g2-g8 (extended seed) matched target RNA paired to guide (above) and relative fraction short g2-g8 matched target bound as a function of hAgo2-guide or EfPiwi-guide concentration. **D**. Release rates (*k*_*off*_) of short g2-g8 target RNA from hAgo2 and EfPiwi guide RNA complexes. **E**. Schematic of short and long g2-g8 matched target RNAs paired to short or long guide RNAs and measured KD values from corresponding EfPiwi-guide complexes. **F**. Relative fraction g2-g8 matched targets (shown in E) bound as a function of EfPiwi-guide concentration.

### Piwi structure facilitates target-pairing downstream of the piRNA seed

Nucleotides immediately downstream of the extended seed, often termed the central region (g9-g12), are not usually used for target recognition by animal miRNAs (Lewis, Shih et al. 2003, Grimson, Farh et al. 2007, Wee, Flores-Jasso et al. 2012, Grosswendt, Filipchyk et al. 2014, Grosswendt, Filipchyk et al. 2014). In hAgo2, central pairing is disfavored by a structural element named the central-gate, which sterically restricts pairing to miRNA nucleotides g9-g12 (Schirle, Sheu-Gruttadauria et al. 2014, Sheu-Gruttadauria, Xiao et al. 2019). Notably, the equivalent gating residues do not extend into the EfPiwi central cleft, but instead extend towards the seed region, making the EfPiwi central cleft wider than that of hAgo2 (**Fig. 4A**). We therefore hypothesized piRNA nucleotides downstream of the seed may have target-binding properties that differ from those of miRNAs.

**Figure 4.**
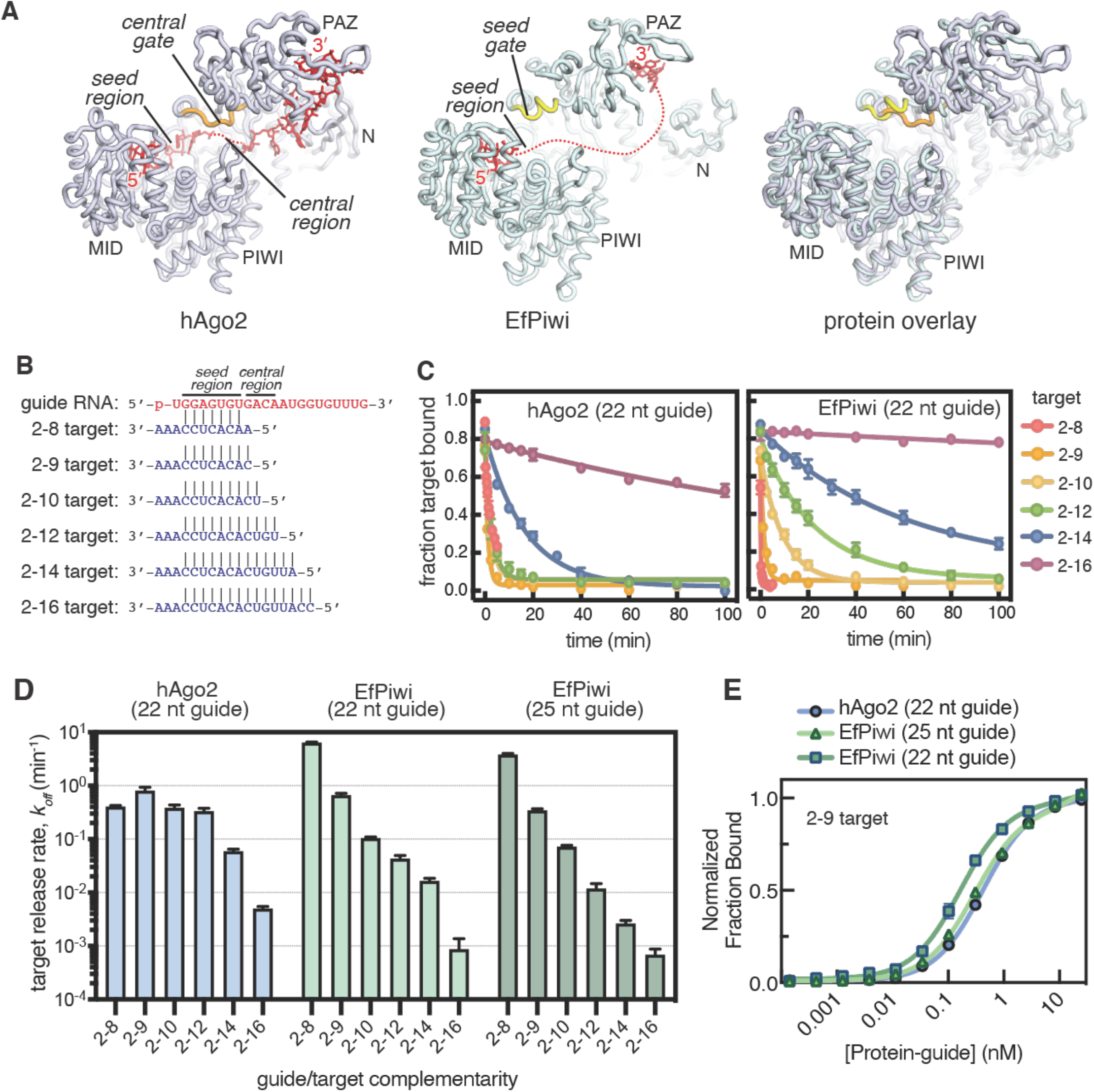
EfPiwi facilitates target pairing downstream of the seed. **A**. Ribbon representation of hAgo2 (left) and EfPiwi (middle) with an overlay (right) showing positions of the central gate and seed gate. Dashed red line indicates unstructured miRNA and piRNA nucleotides. **B**. Schematic of target RNAs of increasing complementarity to guide. **C**. Release of ^32^P-labeled target RNAs from hAgo2-guide (left) or EfPiwi-guide (right) complexes in the presence of excess unlabeled target RNA over time. **D**. Release rates (*k*_*off*_) of target RNAs from hAgo2-guide or EfPiwi-guide complexes (calculated from C and S4A). **E**. Relative fraction 2-9 target RNA bound at equilibrium plotted as a function of hAgo2-guide or EfPiwi-guide concentration.

We measured dissociation of target RNAs of various lengths from EfPiwi or hAgo2 loaded with the same 22 nt guide RNA, and also examined target release from EfPiwi loaded with a 25 nt guide with a 2′-OCH_3_ group on its 3’ nucleotide (**Fig. S4A and B**). hAgo2 released a target complementary to g2-g8 at a rate of 0.81 ± 0.39 min^-1^, and including target complementarity through the central region had little effect on this release rate, consistent with the notion hAgo2 avoids central pairing (**Fig. 4C and D**). In contrast, EfPiwi target release rates decreased with increased guide-target complementarity throughout the central region (**Fig. 4C and D**). Indeed, when using a 25 nt 2′-OCH_3_ guide, the stability of the EfPiwi-piRNA-target ternary complex was directly proportional to the calculated free energy of base pairing through the central region (**Fig. S4C**). We conclude that, unlike Ago2, EfPiwi allows unencumbered guide-target pairing through the central cleft. Notably, a target complementary to g2-g9 bound both EfPiwi and hAgo2 with similar off-rates and affinities (**Fig. 4D and E**). Thus, unfettered guide-target pairing beyond the seed allows EfPiwi to bind targets with affinities equal to or substantially greater than those of hAgo2.

**Figure S4.**
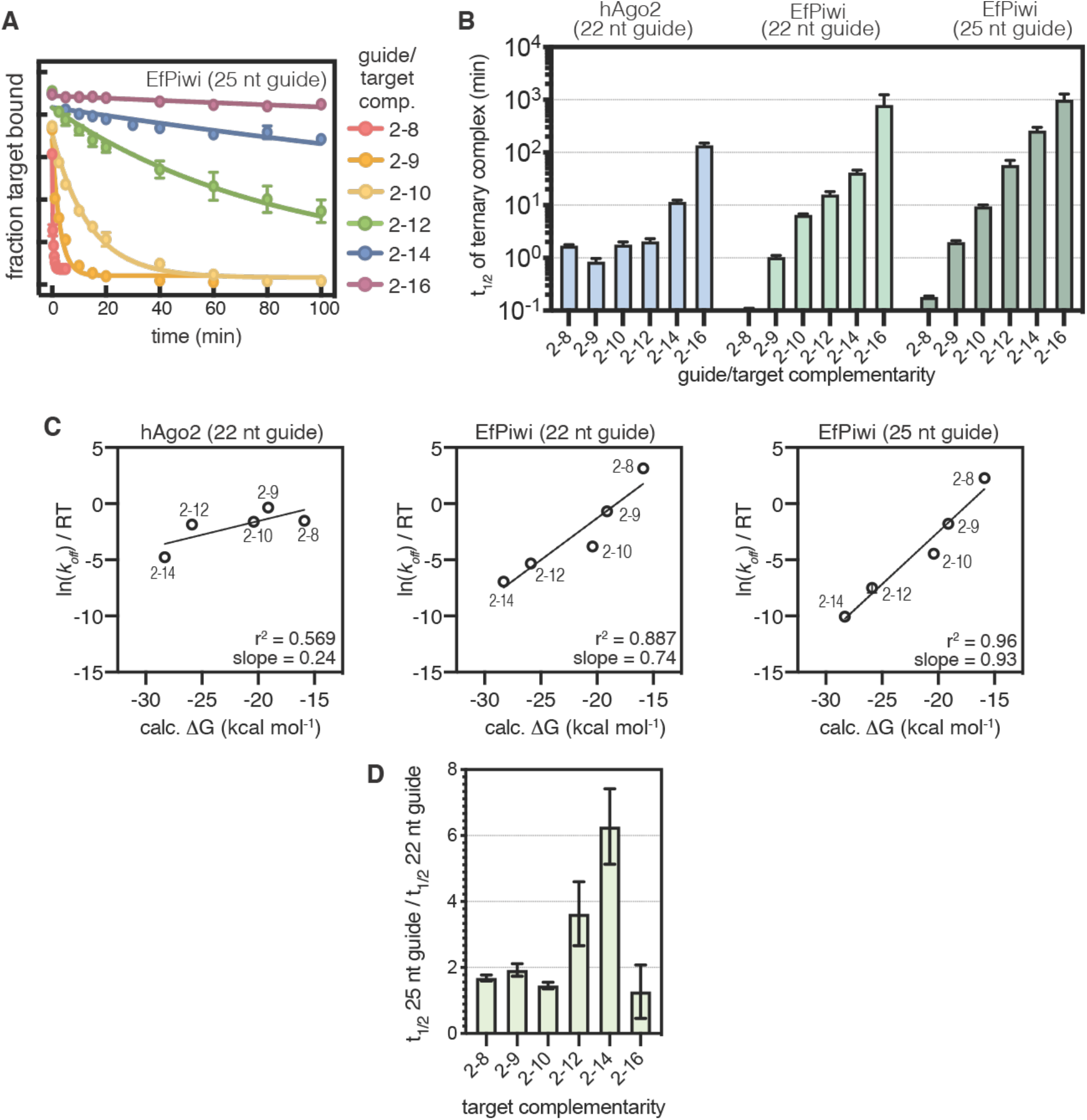
EfPiwi facilitates target pairing downstream of the seed. **A**. Release of target RNAs of increasing complementarity from EfPiwi-25 nt (2’-OCH_3_ on 3’ nucleotide) guide complex. Fraction of input ^32^P-labeled target RNA bound to protein is plotted as a function of time. Target RNA sequences are as shown in Fig. 4B. **B**. Release rates of target RNAs from hAgo2-guide or EfPiwi-guide complexes plotted as half-lives. **C**. Plots of the predicted Gibbs free energy (calc. ΔG) for each piRNA-target duplex versus the natural logarithm of the observed dissociation rates (divided by temperature) from Ago2-guide or EfPiwi-guide complexes. **D**. Bar plot of differences in dissociation rates of targets from EfPiwi loaded with a 22 nt (2’OH) guide or a 25 nt (2’OMe) guide. Differences were only seen in the 2-12 and 2-14 targets, suggesting 3’ end release and piRNA-target wrapping effects.

### Structure of the EfPiwi-piRNA-target complex

To visualize how EfPiwi engages target RNAs we determined a cryo-EM structure of EfPiwi bound to a 24 nucleotide piRNA and a target RNA complementary to piRNA nucleotides g2-g16 to overall resolution of ∼3.5 Å (Fig. 5A). The reconstruction contains a conspicuous 15 bp piRNA-target RNA duplex passing through the EfPiwi central cleft (**Fig. 5B and S5**). Comparison to the binary EfPiwi-piRNA structure reveals that target-binding is associated with widening of the central cleft, which can be roughly described as a ∼25° rotation of the PAZ domain and seed-gate about the L1 stalk (**Fig. 5C**), to accommodate the piRNA-target duplex. A similar conformational change is employed by hAgo2 during target-binding (Schirle, Sheu-Gruttadauria et al. 2014, Sheu-Gruttadauria, Pawlica et al. 2019, Sheu-Gruttadauria, Xiao et al. 2019).

**Figure 5.**
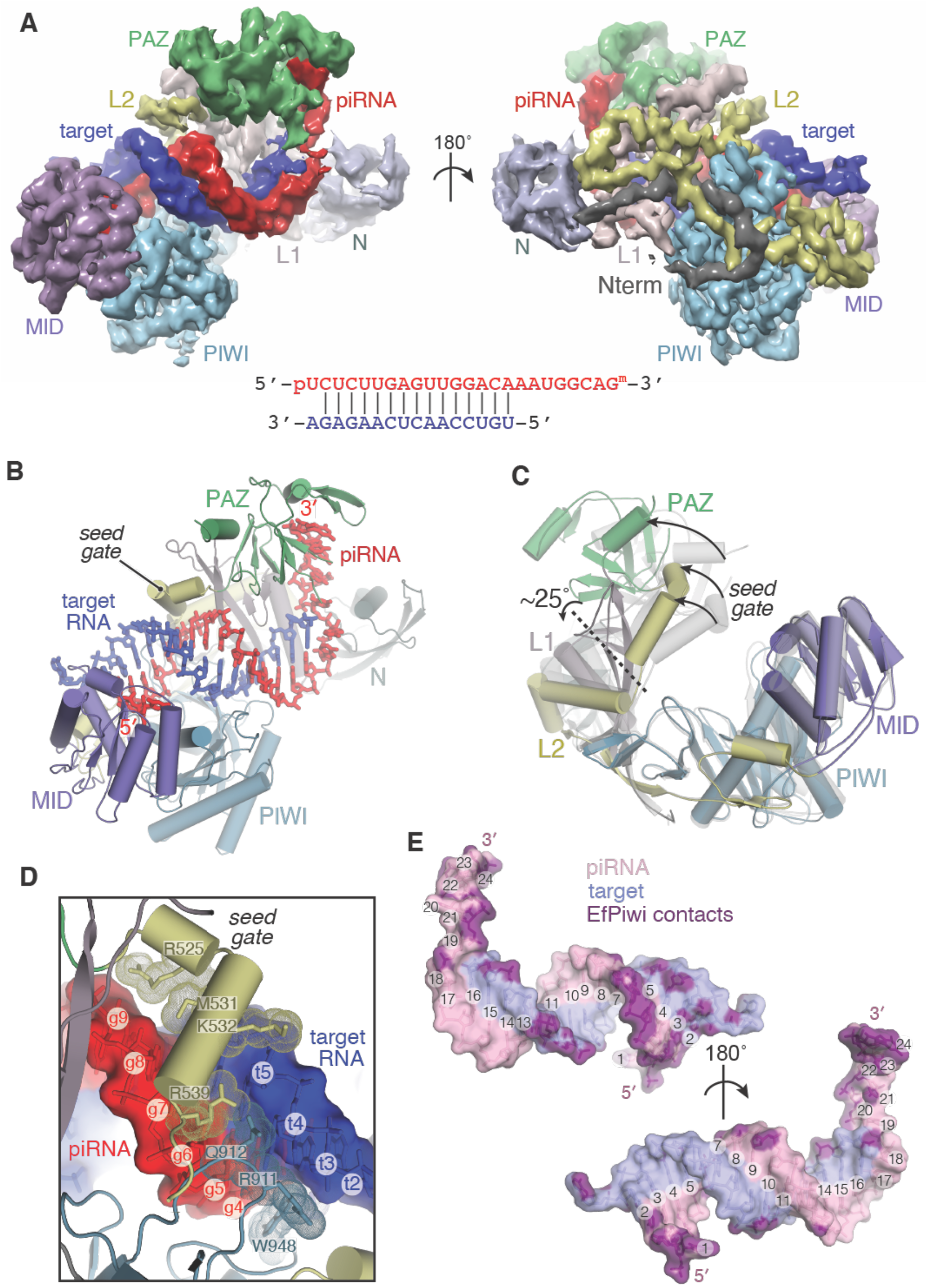
Cryo-EM structure of the EfPiwi-piRNA-target complex. **A**. Segmented reconstruction of EfPiwi-piRNA-target ternary complex with major domains colored distinctly. Guide-target pairing schematic is shown (lower panel). **B**. Cartoon representation of the EfPiwi-piRNA-target ternary complex. **C**. Superposition of EfPiwi-piRNA binary structure (gray) and EfPiwi-piRNA-target ternary complex (colored and solid), highlighting opening of EfPiwi upon target binding. Arrows indicate widening from guide-only to guide-target structures. Dashed line indicates location of primary hinge located in the L1 stalk. **D**. Close up view of interactions between EfPiwi and the minor groove of the piRNA-target duplex. piRNA and target nucleotides numbered at ribose positions. **E**. Surface representation of the piRNA-target duplex with piRNA nucleotides numbered at Watson-Crick face. Non-hydrogen RNA atoms positioned ≤ 4 Å from an EfPiwi atom colored purple.

The 5’ half of the piRNA-target duplex is well-ordered (**Fig. S5F**). EfPiwi contacts the piRNA backbone at positions g1–g7 and g10 and makes hydrophobic and van der Waals interactions with positions 2–9 of the piRNA-target duplex minor groove via aliphatic/aromatic portions of residues from the MID domain (K731), PIWI domain (R911, Q912, W948), and seed-gate (R525, M531, K532, R539) (**Fig. 5D**). The combination of backbone and minor groove contacts may allow EfPiwi to sense deviations from Watson-Crick pairing in this region, consistent with reports that piRNA-targeting *in vivo* is sensitive to mismatches and G:U wobble pairs towards the 5’ half of the piRNA (Goh, Falciatori et al. 2015, Zhang, Kang et al. 2015, Shen, Chen et al. 2018, Zhang, Tu et al. 2018, Halbach, Miesen et al. 2020). hAgo2 similarly probes the miRNA-target minor groove but has a smaller seed-gate and thus only contacts positions 2–7 of the miRNA-target duplex (Schirle, Sheu-Gruttadauria et al. 2014).

Beyond positions 2–9, EfPiwi does not probe the piRNA-target duplex minor groove. Instead, the protein primarily contacts the target (t) RNA sugar-phosphate backbone with numerous residues positioned to make hydrogen bonds/salt linkages to t10–t16 (**Fig. 5E**). By making frequent contacts to the target backbone and ignoring minor grove shape EfPiwi may recognize the overall helical structure of the piRNA-target duplex in a manner that is tolerant of deviations from A-form geometry and thus less sensitive to mismatches and wobble base pairs. Indeed, the density corresponding to base pairs 11-16 of the piRNA-target duplex is not well-resolved, indicative of conformational heterogeneity (**Fig. S5F**). Structural heterogeneity was observed in the density corresponding to piRNA nucleotides g17-g24, but was of sufficient quality to model a single-stranded segment that thread between N and PAZ domains and ends in the 3’-nucleotide binding pocket of the PAZ domain (Tian, Simanshu et al. 2011) (**Fig. S5F**).

**Figure S5.**
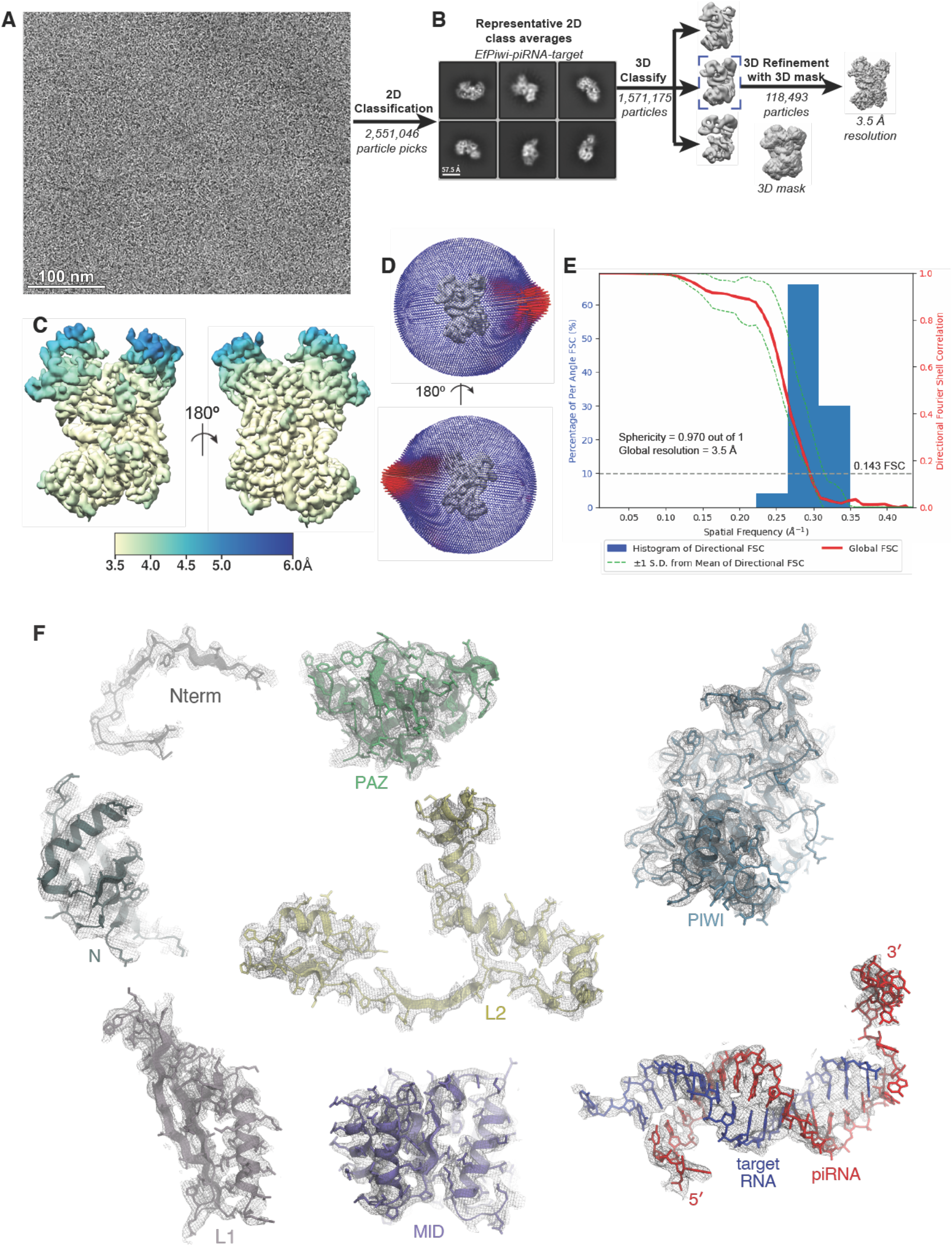
Imaging and processing of the EfPiwi-piRNA-target complex. **A**. Representative cryo-EM micrograph of EfPiwi-piRNA-target complex. **B**. Workflow for processing EfPiwi-piRNA-target complex data set. Particles isolated from micrographs were sorted by reference-free 2D classification. Only particles containing high-resolution features for the intact complex were selected for downstream processing. 3D classification was used to further remove low-resolution or damaged particles, and the remaining particles were refined to obtain a 3.5 Å map. **C**. The EfPiwi-piRNA-target complex map colored by local resolution. **D**. Euler angle distribution plot for the EfPiwi-piRNA-target complex particles. **E**. Directional Fourier Shell Correlation (FSC) plot representing 3D resolution anisotropy in the reconstructed map. Red line shows global FSC; green dashed lines ±1 standard deviation from mean of directional resolutions; blue histograms indicate percentage of directional resolution over the 3D FSC. **F**. EM density quality of EfPiwi-piRNA-target complex. Individual domains of EfPiwi and RNAs fit into the EM density; EM density shown in mesh; protein models shown in cartoon representation (colored as in Fig. 5) with side chains shown as sticks; RNAs shown in stick representation.

### Accurate seed-pairing initiates piRNA target recognition

The ternary EfPiwi-piRNA-target structure suggests EfPiwi can sense piRNA-target mispairing in the seed region but is tolerant of mismatches beyond the seed. We examined this model by measuring the impact of mismatched segments on target recognition (**Fig. 6A and S6A**). The EfPiwi-piRNA complex bound a fully complementary target RNA with an on-rate (*k*_*on*_) of 2.7 ± 0.7 x 10^10^ M^-1^ min^-1^. This *k*_*on*_ closely matches values reported for mouse Ago2 (Salomon, Jolly et al. 2015) and hAgo2 (Klum, Chandradoss et al. 2018), revealing Piwi and Ago proteins bind targets at similar rates. Introducing 3 nt mismatched segments opposite the piRNA seed reduced *k*_*on*_ by more than two orders of magnitude (**Fig. 6B**). In contrast, mismatched segments beyond the seed had only minor effects (≤ 2.5-fold) on target binding rate. These results indicate EfPiwi relies almost exclusively on accurate seed-pairing for initiating target recognition, explaining why seed-complementarity is essential for piRNA targeting *in vivo* (Goh, Falciatori et al. 2015, Shen, Chen et al. 2018, Zhang, Tu et al. 2018, Halbach, Miesen et al. 2020).

**Figure 6.**
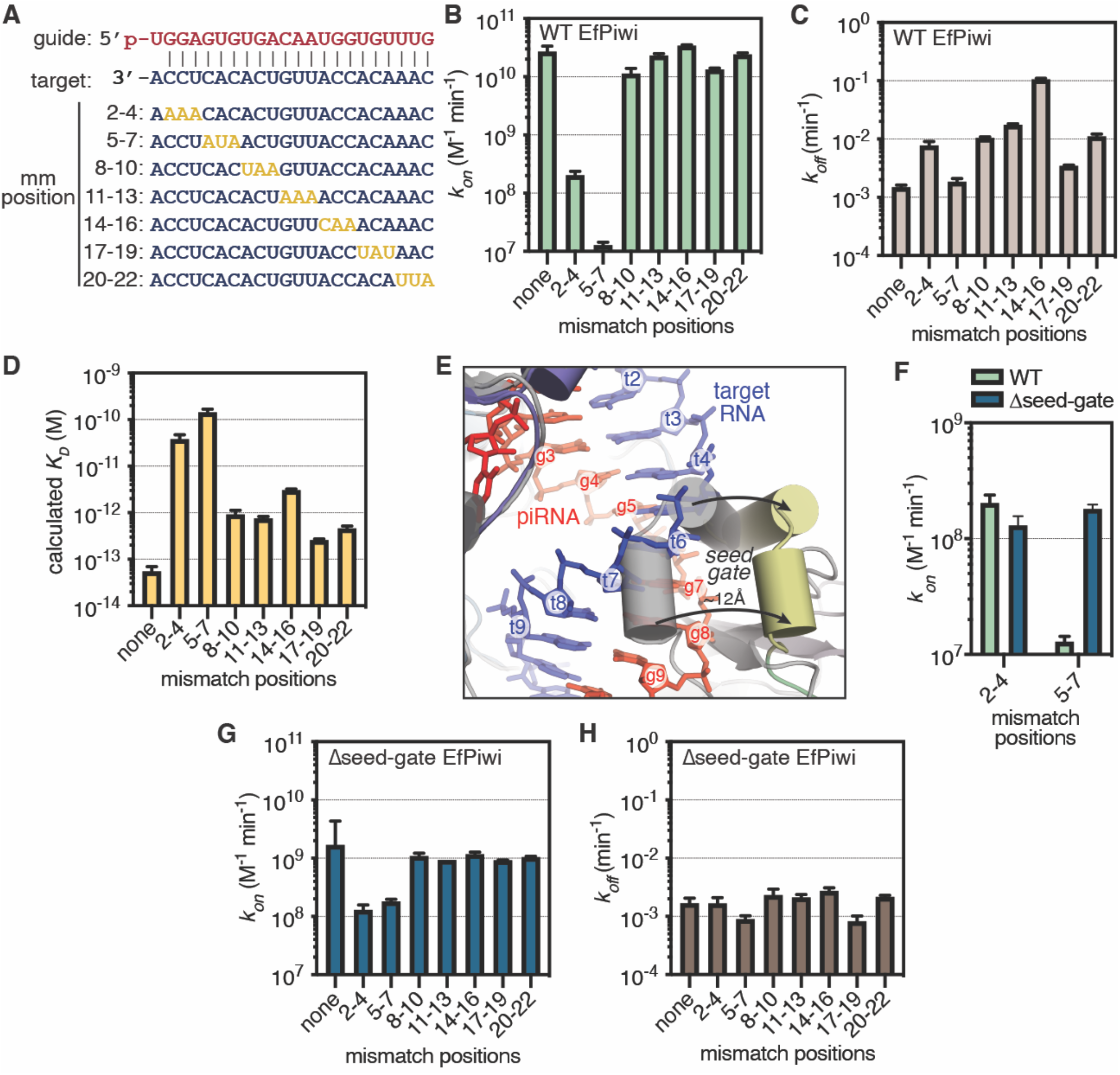
piRNA-targeting with mismatches and role of the seed-gate. **A**. Guide-target pairing schematic for mismatched targets. Mismatches colored gold. **B**. Association rates of ^32^P-labeled target RNAs with three consecutive mismatches to wild-type EfPiwi. **C**. Dissociation rates of ^32^P-labeled target RNAs with three consecutive mismatches from wild-type EfPiwi. **D**. Calculated dissociation constants (*K*_*D*_) for target RNAs. **E**. Superposition of EfPiwi-piRNA binary structure (gray and semitransparent) and EfPiwi-piRNA-target ternary complex (colored), showing that the seed gate opens ∼12 Å upon target binding. Arrows indicate direction of movement from guide-only to target-bound structures. **F**. Association rates of ^32^P-labeled target RNAs with mismatches to g2-g4 (5’ end of seed) and g5-g7 (3’ end of seed) to the wild-type EfPiwi and Δseed-gate EfPiwi. **G**. Association rates of ^32^P-labeled target RNAs with three consecutive mismatches to Δseed-gate EfPiwi. **H**. Dissociation rates of ^32^P-labeled target RNAs with three consecutive mismatches from Δseed-gate EfPiwi.

We also measured the effects of mismatches on target release rates (*k*_*off*_) (**Fig. 6C and S6B**). Release of the fully complementary target was very slow, with a half time (t_1/2_) of > 8 hours. Mismatched segments generally had mild to moderate effects on target release, increasing *k*_*off*_ 2–12-fold. One exception was a target with mismatches at g14-g16, which was released ∼70-fold faster than the full complementary target, resembling targets with pairing limited to the seed and central regions (**Fig. 4D**). Taken with the structure of the ternary complex, we suggest that g14-g16 pairing may be particularly important for driving sustained release of the piRNA 3’ end, which is necessary for piRNA-target wrapping and duplex propagation beyond the piRNA central region.

Dissociation constants (*K*_*D*_) for extensively paired targets were too small to measure. However, *K*_*D*_ values could be calculated from *k*_*off*_ / *k*_*on*_ ratios, which indicate EfPiwi can bind all targets with high (sub-nM) affinity in our purified system, if given a sufficient amount of time (**Fig. 6D**). Affinity is most sensitive to mismatches in the piRNA seed region due to low on-rates. Affinity is also sensitive to g14-g16 mismatches due to higher target release rates, which may relate to the finding that piRNA targeting in *C. elegans* is disrupted by mismatches surrounding g15-g18 (Shen, Chen et al. 2018).

**Figure S6.**
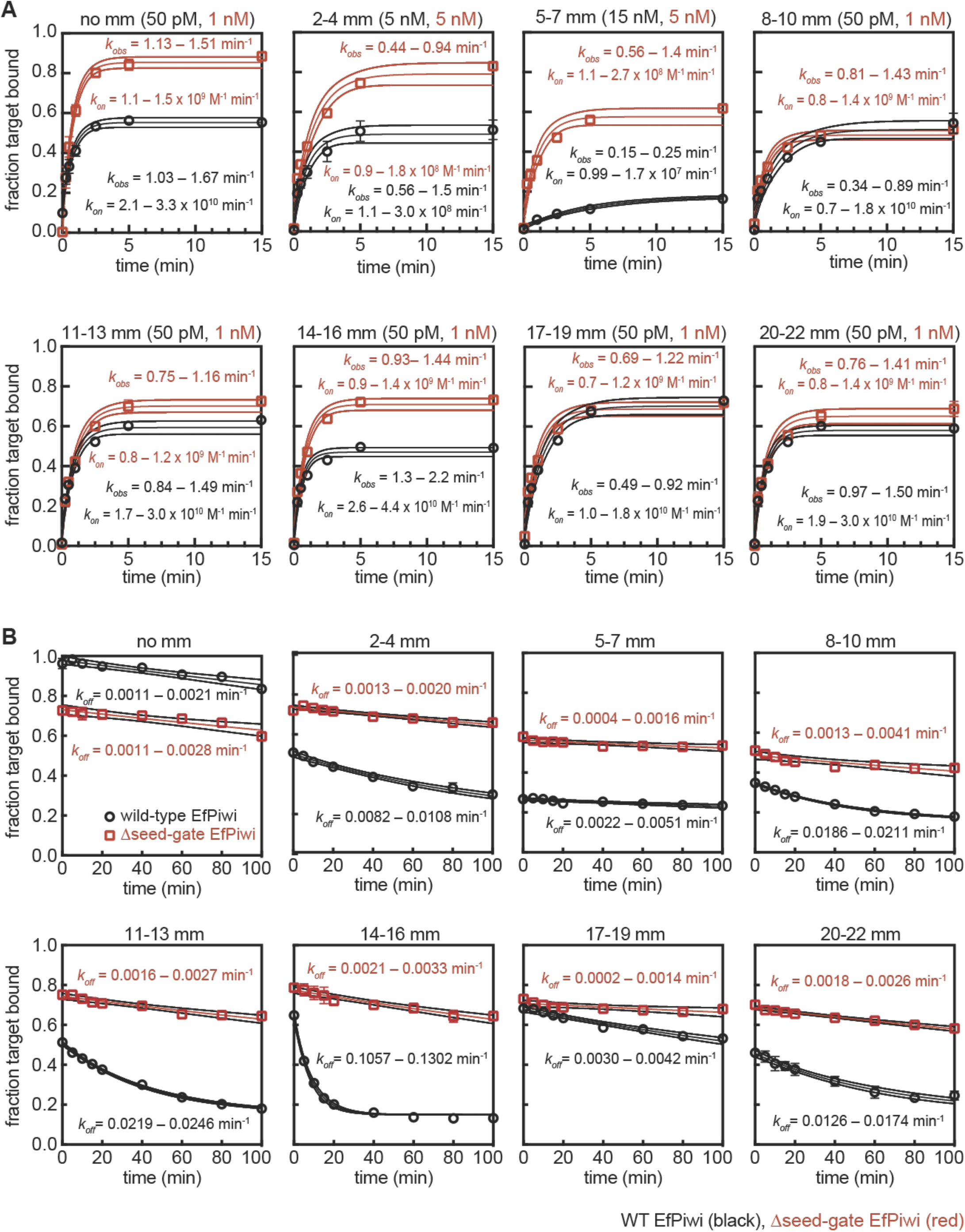
EfPiwi target binding data. **A**. Raw data for *k*_*on*_ values shown in Fig. 6B, 6F and 6G. Plots of target RNAs with mismatches (sequences shown in Fig. 6A) binding to EfPiwi-piRNA complexes over time. Protein concentrations used in each experiment are indicated at top of each graph. 95% confidence limits of observed association rates (*k*_*obs*_) and *k*_*on*_ values indicated. WT EfPiwi (black), Δseed-gate EfPiwi (red). **B**. Raw data for *k*_*off*_ values shown in Fig. 6C and 6H. Plots of target RNAs with mismatches (mm) dissociating from EfPiwi-piRNA complexes over time. All data were fit to a plateau value of 0.15. 95% confidence limits of *k*_*on*_ values indicated. All data points were measured three times. Error bars indicate SEM. Center line indicates best fit to data. Surrounding lines indicate 95% confidence limits.

### The seed-gate regulates propagation of piRNA-target pairing

EfPiwi target-binding is exceptionally sensitive to g5-g7 mismatches, which reduced *k*_*on*_ > 2000-fold (**Fig. 6B**). This was surprising because Ago2 *k*_*on*_ values are relatively insensitive to mismatches towards the 3’ end of the seed (Chandradoss, Schirle et al. 2015, Salomon, Jolly et al. 2015). We sought a structural explanation for the unique targeting behavior of EfPiwi. Comparing EfPiwi-piRNA and EfPiwi-piRNA-target structures shows pairing to the 3’ end of the seed requires a ∼12 Å movement of the seed-gate to avoid severe steric clashes with base pairs 5-8 of the piRNA-target duplex (**Fig. 6E**). Taken with the observation that, upon opening, the seed-gate makes multiple contacts to the piRNA-target duplex minor grove (**Fig. 5D**), we speculated the seed-gate might be a sensor for accurate piRNA-target pairing in the 3’ end of the seed. To explore this idea, we generated a Δseed-gate EfPiwi mutant, in which the seed gate (residues L520-H537) was replaced with a Gly_6_ linker. The Δseed-gate EfPiwi bound the g2-g4 mismatched target at the same rate as wild type EfPiwi (**Fig 6F**). In contrast, seed-gate removal increased *k*_*on*_ for the g5-g7 mismatched target 14-fold (**Fig. 6F**). We propose EfPiwi uses the seed-gate as a physical barrier to inhibit propagation of the piRNA-target duplex the absence of faithful base pairing to the 3’ end of the seed.

We also examined binding of Δseed-gate EfPiwi to targets with mismatches beyond the seed region. All targets with intact seed-complementarity, including the fully complementary target, were bound by Δseed-gate EfPiwi ≥ 10-fold more slowly than wild type EfPiwi (**Fig. 6G**). This observation indicates the seed-gate also facilitates propagation of piRNA-target duplexes established through transient interactions with 5’ end of the seed. A similar function was proposed for the structural equivalent of the seed-gate in hAgo2A (termed helix-7 in hAgo2) (Klum, Chandradoss et al. 2018). Finally, Δseed-gate EfPiwi released all mismatched target RNAs more slowly than wild type EfPiwi, indicating a general role in breaking piRNA-target pairing (**Fig. 6H**) as also suggested for helix-7 in hAgo2 (Klum, Chandradoss et al. 2018). We conclude that the seed-gate enables fidelity in piRNA-targeting by both facilitating extension of piRNA-target duplexes initiating at the seed 5’ end and blocking propagation of piRNA-target duplexes lacking sufficient complementary to the seed 3’ end.

### A manganese cofactor stimulates target cleavage by EfPiwi

Once bound, PIWI proteins often endonucleolytically cleave target RNAs (Kawaoka, Hayashi et al. 2009, De Fazio, Bartonicek et al. 2011, Reuter, Berninger et al. 2011, Xiol, Spinelli et al. 2014, Goh, Falciatori et al. 2015, Nishida, Iwasaki et al. 2015, Matsumoto, Nishimasu et al. 2016, Yamaguchi, Oe et al. 2020). We examined cleavage of a target RNA fully complementary to the piRNA bound to EfPiwi. In initial experiments, cleavage by EfPiwi was nearly undetectable, leading us to speculate a cofactor may be missing. Because Mg^2+^ is the only known cofactor required for target cleavage by Argonaute (Schwarz, Tomari et al. 2004, Rivas, Tolia et al. 2005), we screened other divalent cations and found cleavage by EfPiwi is specifically stimulated by Mn^2+^ (**Fig. 7A, left**). In contrast, hAgo2 endonuclease activity was supported by most divalent cations tested (**Fig. 7A, right**). Thus, the divalent cation requirement for EfPiwi is distinct from hAgo2, and instead similar to many prokaryotic Ago proteins, which are also more active in the presence of Mn^2+^ than Mg^2+^ (Yuan, Pei et al. 2005, Wang, Juranek et al. 2009, Swarts, Hegge et al. 2015, Kaya, Doxzen et al. 2016). Physiological Mn^2+^ concentrations (0.1 mM) supported weak cleavage activity, which was further stimulated by 1 mM Mg^2+^ (**Fig. S7C**). Thus, EfPiwi may employ Mn^2+^ in combination with Mg^2+^ to catalyze target cleavage *in vivo*.

**Figure 7.**
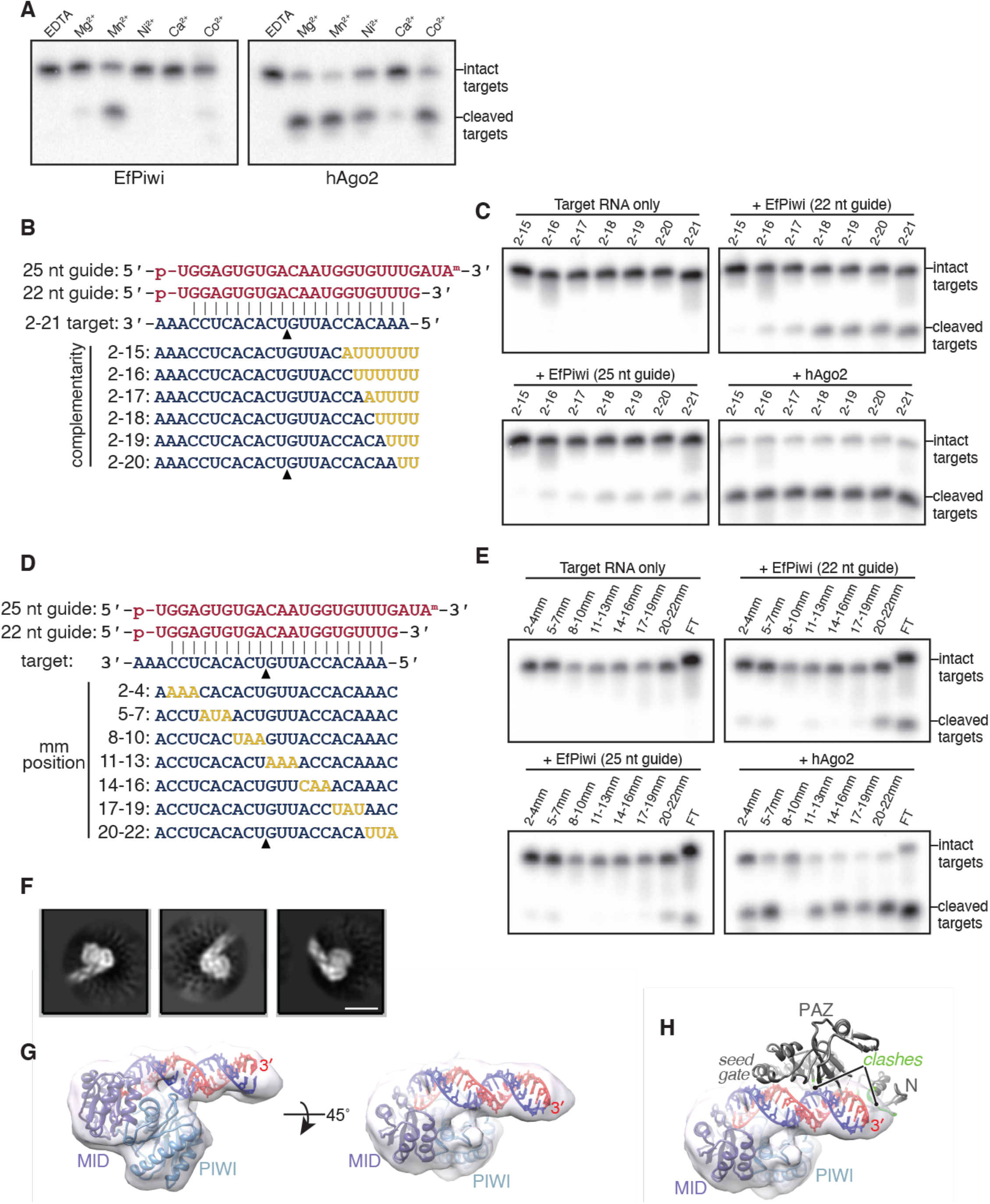
Target cleavage by EfPiwi requires a manganese cofactor and is driven by extended piRNA-target pairing. **A**. Cleavage of g2-g21 matched ^32^P-labeled target RNA by hAgo2 or EfPiwi in the presence of various divalent cations (2 mM each). **B**. Guide-target pairing schematic for targets with mismatches shown in gold. Arrowhead indicates site of cleavage. **C**. Denaturing polyacrylamide gel electrophoresis showing cleavage of ^32^P-labeled target RNAs of increasing complementarity by EfPiwi loaded with a 22 or 25 nt guide or hAgo2 loaded with a 22 nt guide. **D**. Guide-target pairing schematic for targets with mismatches shown in gold. Arrowhead indicates site of cleavage. **E**. Denaturing polyacrylamide gel electrophoresis showing cleavage of ^32^P-labeled target RNAs with three consecutive mismatches by EfPiwi loaded with a 22 or 25 nt guide or hAgo2 loaded with a 22 nt guide. **F**. Two-dimensional class averages of EfPiwi-piRNA complex bound to a target with complementarity from g2-g25. Scale bar is 5 nm. **G**. Three-dimensional cryo-EM reconstruction of the EfPiwi-guide complex bound to a target paired from g2-g25 fit with the MID/PIWI lobe and an extended piRNA-target duplex. piRNA 3’ end indicated. **H**. Docking the EfPiwi-guide complex bound to a target paired from g2-g16 onto the MID/PIWI domains of the g2-g25 target-bound model reveals steric clashes (green) between the EfPiwi N/PAZ lobe and the extended piRNA-target duplex.

**Figure S7.**
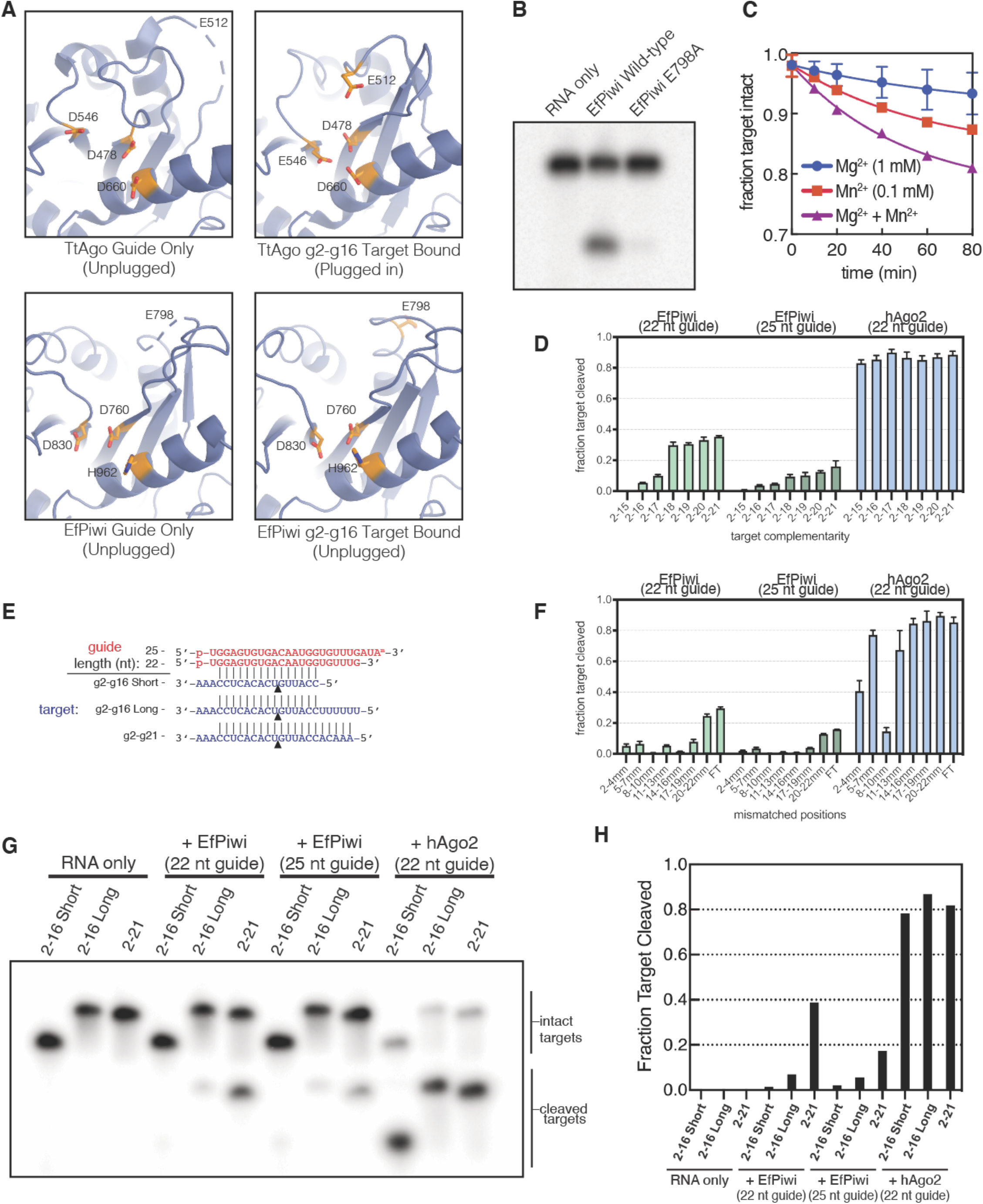
EfPiwi target cleavage employs a “plugged in” catalytic tetrad and is moderately affected by guide length. **A**. TtAgo (upper left, PDB 3DLH) and EfPiwi (lower left) adopt an inactive, or “unplugged” conformation with no target bound, with E512 or E798, respectively, away from the active site. EfPiwi with a g2-g16 paired target bound (lower right) retains the inactive conformation, whereas TtAgo bound to a target of the same complementarity (upper right, PDB 3HJF) is in a catalytically active conformation. **B**. Denaturing polyacrylamide gel electrophoresis shows mutation of E798 to alanine greatly diminishes target cleavage by EfPiwi. **C**. Time course showing cleavage of g2-g21 paired ^32^P-labeled target RNA by EfPiwi in the presence of Mg^2+^, Mn^2+^, or both at approximate physiological divalent cation concentrations. **D**. Quantification of cleavage of ^32^P-labeled target RNAs of increasing complementarity by EfPiwi loaded with a 22 or 25 nt guide or hAgo2 loaded with a 22 nt guide shown in Fig 7C. Plotted values are means of at least three replicate experiments. Error bars indicate SEM. **E**. Schematic of target pairing of short and long targets to EfPiwi loaded with a 22 or 25 nt guide. **F**. Quantification of cleavage of ^32^P-labeled target RNAs with three consecutive mismatches by EfPiwi loaded with a 22 or 25 nt guide or hAgo2 loaded with a 22 nt guide shown in Fig 7E. Plotted values are means of at least three replicate experiments. Error bars indicate SEM. **G, H**. Denaturing polyacrylamide gel electrophoresis and quantification showing cleavage of ^32^P-labeled target RNAs by EfPiwi loaded with a 22 or 25 nt guide or hAgo2.

### Extended piRNA-target pairing stimulates target cleavage

The target cleavage mechanism is best understood in *T. thermophilus* Ago (TtAgo). Binding targets with complementarity extending to g15 induces TtAgo to shift from a catalytically inactive (“unplugged”) to an active (“plugged in”) conformation (Sheng, Zhao et al. 2014). This conformational change involves movement of catalytic residue E512 into the endonuclease active site. In contrast to TtAgo, our EfPiwi structure shows a piRNA-target duplex extending to g16 does not induce E798 (the equivalent to TtAgo E512) to shift into the active site (**Fig S7A**). We hypothesized EfPiwi may either employ a distinct catalytic mechanism or require further target pairing to stimulate cleavage.

We first examined target cleavage by an EfPiwi mutant in which E798 was changed to alanine, which almost completely abolished generation of a cleavage product (**Fig. S7B**). Taken with the previous finding that the analogous E708A mutation inhibited cleavage by Siwi (Matsumoto, Nishimasu et al. 2016), this result supports the hypothesis that, like TtAgo, PIWI proteins undergo an “unplugged” to “plugged-in” transition during catalysis (Matsumoto, Nishimasu et al. 2016). We next examined the extent of piRNA-target complementarity required for cleavage. Cleavage of a target with complementarity limited to g2-g15 was almost undetectable, even when treating 1 nM target with a vast excess (100 nM) of EfPiwi-guide complex for one hour (**Fig. 7C, S7D, and S7E-F**). Cleavage activity increased with greater complementarity towards the guide RNA 3’ end, reaching a plateau with pairing to g2–g18 (**Fig. 7B-C, and S7D**). For comparison, the same concentration of hAgo2-guide complex effectively cleaved all targets tested (**Fig. 7B-C and S7D**). Finally, we examined how mismatched segments in the guide-target duplex influence cleavage activity. Mismatched segments (3-nts in length) expected to occur within the EfPiwi central cleft reduced target cleavage. In contrast, g20-g22 mismatches at the end of the piRNA-target duplex did not. Mismatched segments expected to reside between the seed-gate and N-domain (g8-g10 – g17-g19) were especially detrimental, reducing cleavage to near background levels (**Fig. 7D-E and S7F**).

### Extended piRNA-target pairing fully opens the EfPiwi central cleft

To gain insight into why target cleavage requires extended piRNA-target pairing, we examined the structure of the EfPiwi-piRNA complex bound to a target RNA with complementarity to g2-g25 by cryo-EM. While the piRNA-target duplex was obvious in 2D averages, the EfPiwi protein appeared smaller than expected (**Fig. 7F**). Indeed, a 3D reconstruction of this complex determined to ∼7.0 Å resolution was consistent with the MID and PIWI domains bound to the end of the piRNA-target duplex (**Fig. 7G**). These results indicate that, upon extended pairing, the piRNA-target duplex remains anchored to the MID-PIWI lobe while forcing open the N-PAZ lobe, which becomes disordered relative to the rest of the complex. Supporting this idea, docking the EfPiwi-piRNA-target model (with pairing restricted to g2-g16) onto the reconstruction with extended target pairing reveals steric clashes between the RNA and loops in the L2 and N domains (**Fig. 7H**).

We suggest that, unlike TtAgo, target cleavage by EfPiwi requires substantial widening of the central cleft, driven by formation of a piRNA-target duplex extending beyond the N domain. Mismatched segments ≥ 3 nt in length impart flexibility to the piRNA-target duplex (Sheu-Gruttadauria, Pawlica et al. 2019), leading to potential misalignment of the scissile phosphate and a reduced propensity to drive open the central cleft, both of which we propose would inhibit target cleavage. Using this mechanism, EfPiwi creates a target-cleavage recognition site larger than that of TtAgo and hAgo2, and thus may avoid cleavage of targets lacking extensive piRNA-complementary. Notably, piRNA cleavage targets identified in mouse testes are generally free of RNA bulges and mismatched segments ≥3 nt in length (Goh, Falciatori et al. 2015, Zhang, Kang et al. 2015, Wu, Fu et al. 2020), and target cleavage by immuno-purified Miwi also required extended piRNA-target complementarity (Reuter, Berninger et al. 2011). We suggest central cleft widening driven by extended piRNA-target pairing may be a feature of endonucleolytic mechanisms used by diverse PIWI proteins.

## Discussion

piRNAs and miRNAs are genetic regulators found in nearly all animal species, and are suggested to have helped to usher in the era of multicellular animal life (Grimson, Srivastava et al. 2008). Our analysis of EfPiwi structure and comparison to hAgo2 reveals PIWI and AGO clade proteins possess divergent functional properties, enabling miRNAs and piRNAs to carry out the distinct roles in animal biology.

The primary function of miRNAs is to repress cellular mRNAs in order to orchestrate development and maintain homeostasis. Thus, miRNAs and their targets evolve together to favor recognition. The AGO targeting mechanism supports this relationship: hAgo2 creates a strong seed, which is generally sufficient for target recognition, and uses its central gate to discourage miRNA-target pairing between seed and supplementary regions. Thus, miRNA-recognition elements are short and well-defined, allowing each miRNA to regulate hundreds of targets, and enabling straightforward acquisition and loss of miRNA-recognition sites over evolutionary time (Nozawa, Fujimi et al. 2016, Simkin, Geissler et al. 2020).

piRNAs can also regulate cellular RNAs, but the primary piRNA function is silencing TEs, parasitic genetic material under selective pressure to escape recognition. It has been proposed that flexibility in target recognition may enable piRNAs to respond to evolving TEs (Brennecke, Aravin et al. 2007). On the other hand, considering the immense sequence diversity of the piRNA repertoire, targeting must also be stringent enough to avoid inadvertently silencing cellular RNAs. EfPiwi accomplishes this balancing act by creating a weak seed, making target recognition dependent on downstream pairing. Seed-pairing stringency is nonetheless achieved by pre-organizing the seed 5’ end to accelerate interactions with target sites, and by placement of the seed-gate, which requires 3’ seed pairing to open and enable propagation of the piRNA-target duplex. Targeting flexibility, in terms of binding, is provided beyond the seed-gate, were the central cleft allows unencumbered piRNA-target pairing and is generally tolerant of helical imperfections arising from piRNA-target mismatches. Stringency is further established for target cleavage by the requirement of a rigid piRNA-target duplex long enough to push beyond the central cleft and activate the endonuclease mechanism. We suggest the combined processes make piRNA-targeting specific enough to minimize spurious off-targeting yet flexible enough to allow recognition of transcripts between related TE families, and to limit TE escape from piRNA surveillance through mutation.

Finally, our results provide insights for considering silencing mechanisms downstream of piRNA target recognition. We note that EfPiwi target cleavage is far less efficient than that of hAgo2, with the majority of target molecules bound but not cleaved by EfPiwi after extended amounts of time. This raises the possibility that our *in vitro* cleavage conditions might lack cellular factors that stimulate and/or regulate EfPiwi activity *in vivo*. Alternatively, cellular targets may often be bound by EfPiwi in cells without ever being cleaved. Indeed, high-affinity target binding (**Fig. 6A-D**) requires less piRNA-target complementarity than even nominal cleavage observed in vitro (**Fig. 7D-E**). We also note that the release of targets bound by the Piwi-piRNA complex is very slow, on par with the lifetimes of the most stable mRNAs in ES cells (Herzog, Reichholf et al. 2017). Thus, once Piwi engages a target it could potentially remain bound for the remainder of the transcript’s existence. Considering the diversity and abundance of the cellular piRNA pool, we suggest target transcripts may accumulate numerous bound Piwi molecules, creating multivalent assemblies in which Piwi proteins act in concert to recruit of histone/DNA methylation factors to target loci in the nucleus, as well as traffic cytoplasmic target RNAs into phase-separated compartments associated with silencing and piRNA production (Nott, Petsalaki et al. 2015).

## Acknowledgments

We are grateful to Noriko Funayama for *Ephydatia fluviatilis piwi-a* and *piwi-b* cDNA clones and to Irwin H. Segel for advice about measuring binding reactions with very slow off-rates. Research of G.C.L. is supported by NIH grant R21AG067594 and an Amgen Young Investigator Award. Research of I.J.M. is supported by NIH grant R35GM127090.

## Author Contributions

T.A. prepared EfPiwi-piRNA and hAgo2-miRNA samples, performed biochemical experiments, built EfPiwi models, and co-wrote the manuscript. S.C. prepared cryo-EM samples, collected data, produced high resolution reconstructions, and assisted T.A. with model building. S.M.H. identified and developed EfPiwi as a source of active PIWI protein. Y.X. helped develop EfPiwi and establish purification protocols. G.C.L. provided structural insights and guidance in cryo-EM data collection and analysis. I.J.M. provided structural and mechanistic insights and co-wrote the manuscript.

## Data availability

Maps for the EfPiwi-piRNA and EfPiwi-piRNA-target complexes were deposited in the Electron Microscopy Data Bank under accession IDs EMD-23061 and EMD-23063, respectively. Corresponding atomic models were deposited in the Protein Data Bank under accession IDs 7KX7 and 7KX9. The EfPiwi(MID/PIWI)-piRNA-long-target complex map was deposited in the Electron Microscopy Data Bank under accession ID EMD-23062.

## EXPERIMENTAL MODEL AND SUBJECT DETAILS

### Bacterial strains and plasmids

Bacteria used for cloning were chemically competent *E. coli* OmniMAX™ (Thermo Fisher). Bacteria used for production of bacmid were DH10Bac™ chemically competent *E. coli* (Thermo Fisher). Plasmids used are listed in the Key Resources table.

### Bacterial media and growth conditions

All bacterial cultures were grown in Luria-Bertani (LB) medium at 37 °C. When needed, media was supplemented with one or more of the following antibiotics at the following concentrations: ampicillin (100 μg/mL), kanamycin (40 μg/mL), tetracycline (5 μg/mL), gentamycin (7 μg/mL), 5-Bromo-4-Chloro-3-Indolyl β-D-Galactopyranoside (X-gal, 20 μg/mL in dimethylformamide), and/or Isopropyl β-D-1-thiogalactopyranoside (IPTG, 1 mM).

### Insect cell media and growth conditions

Sf9 cells were grown in Lonza Insect XPRESS™ medium supplemented with penicillin (100 units/mL), streptomycin (100 μg/mL), and L-glutamine (2.92 mg/mL) in suspension at 27 °C.

## METHOD DETAILS

### Cloning and mutagenesis

DNA fragments encoding truncations EfPiwi of were generated by PCR using a cDNA clone encoding *Ephydatia fluviatilis* Piwi A (generously provided by Noriko Funayama, Kyoto University) as template. PCR products were cloned as SfoI-XhoI fragments into a modified form of pFastBac HTA (Thermo Fisher) to generate expression plasmids for the Bac-to-Bac baculovirus expression system (Thermo Fisher). Mutations were made using the wild-type EfPiwi expression plasmid as a template and amplifying the protein using oligonucleotide primers containing the desired mutation. The methylated parent plasmid was degraded with DpnI (New England Biolabs), and newly generated plasmid was transformed into OmniMAX chemically competent *E. coli* (Thermo Fisher). These plasmids were used in the Bac-to-bac expression system. All constructs and mutations were confirmed by Sanger sequencing (Genewiz).

### Preparation of hAgo2-guide complexes

Human Ago2 proteins loaded with guide-1 were purified as described previously (Schirle, Sheu-Gruttadauria et al. 2014). His-tagged hAgo2 was purified from Sf9 cells using a baculovirus system (Thermo Fisher). Cells were lysed in Lysis Buffer (300mM NaCl, 0.5 mM TCEP, 50 mM Tris, pH 8) using a single pass through a M-110P lab homogenizer (Microfluidics). Lysate was cleared by centrifugation and the soluble fraction was applied to Ni-NTA resin (Qiagen) and incubated for 1.5 h. The resin was washed with Nickel Wash Buffer (300 mM NaCl, 300 mM imidazole, 0.5 mM TCEP, 50 mM Tris, pH 8). Co-purifying cellular RNAs were degraded with micrococcal nuclease (TakaraBio) on-resin in Nickel Wash Buffer supplemented with 5 mM CaCl_2_. The resin was washed again with Nickel Wash Buffer and then eluted in four column volumes of Nickel Elution Buffer (300 mM NaCl, 300 mM imidazole, 0.5 mM TCEP, 50 mM Tris, pH 8). Eluted hAgo2 was incubated with synthetic miRNA concurrent with overnight cleavage of the N-terminal His_6_ tags using TEV protease and dialysis against Hi-Trap Dialysis Buffer (300 mM NaCl, 15 mM imidazole, 0.5 mM TCEP, 50 mM Tris, pH 8) at 4 °C. The dialyzed protein was then passed through a Hi-Trap Chelating column (GE Healthcare) and the unbound material was collected. The hAgo2 molecules loaded with the desired miRNA were purified using a modified Arpon method (Flores-Jasso, Salomon et al. 2013). Loaded molecules were further purified by size exclusion using a Superdex 200 Increase 10/300 column (GE Healthcare) in High Salt SEC Buffer (1M NaCl, 0.5mM TCEP, 50mM Tris pH 8). The final protein was dialyzed against low salt buffer (10 mM Tris pH 8, 100 mM NaCl, 0.5 mM TCEP), concentrated, aliquoted, flash frozen and stored at -80 °C. Samples were thawed slowly on ice for all subsequent experiments. Concentration of hAgo2 complex was determined by absorbance at λ=280 nm (extinction coefficient 74,720 M^-1^ cm^-1^).

### Preparation of EfPiwi-guide complexes

*Ephydatia fluviatilis* Piwi A (EfPiwi) proteins loaded with the guide RNA were purified using an adapted version of the hAgo2 protocol. EfPiwi protein was expressed as a His_6_-Strep-Strep-TEV-EfPiwi construct (HSST-EfPiwi). EfPiwi protein was expressed using the Bac-to-Bac Baculovirus Expression System (Thermo Fisher) and Sf9 cells. Sf9 infection with EfPiwi-expressing baculovirus was set up in 750 ml cultures of 1275 x 10^6^ cells and infection was incubated for 72 hours at 27 °C. Each 750 ml culture of Sf9 cells was pelleted and resuspended in 25 ml lysis buffer (50 mM NaH_2_PO_4_ pH 8, 300 mM NaCl, 0.5% Triton-X 100, 5% glycerol, and 0.5 mM TCEP) and lysed using Dounce homogenization. The debris was pelleted by centrifugation, and clarified lysate was incubated with 1 ml packed Ni-NTA resin (QIAGEN) per 750 ml culture for 1 hour at 4°C, followed by washing twice in Nickel Wash Buffer (50 mM Tris pH8, 300 mM NaCl, 20 mM imidazole, 0.5 mM TCEP). Following this, resin was washed once using wash buffer supplemented with 5 mM CaCl_2_ in preparation for micrococcal nuclease treatment to degrade co-purifying cellular RNAs. Washed resin was resuspended in Nickel Wash Buffer supplemented with 5 mM CaCl_2_ to a final volume of 20 ml, and 5 μl micrococcal nuclease (TakaraBio) was added per 750 ml culture and this was incubated at room temperature for 1 hour, inverting gently every 15 minutes to resuspend resin. Following 3 more washes in Nickel Wash Buffer without CaCl_2_, protein was eluted with 6 column volumes of Nickel Elution Buffer (Wash Buffer supplemented with 300 mM imidazole). Eluted protein was supplemented with 5 mM EGTA to remove any remaining calcium, and 7 nmol guide RNA (Integrated DNA Technologies) per 750 ml Sf9 culture was added. This was dialyzed overnight against 50 mM Tris pH8, 300 mM NaCl, 0.5 mM TCEP buffer along with TEV protease to cleave the His_6_-Strep-Strep tag.

After dialysis, the buffer was supplemented with 30 mM imidazole and an additional 100 mM NaCl and passed through a 1 ml Hi-Trap Chelating column (GE Healthcare) to capture the N-terminal fragment that was cleaved by the TEV protease. EfPiwi protein in the flow through was supplemented with 0.02% CHAPS and 2 mM MgOAc in preparation for guide-specific capture.

To prepare a resin for capturing guide-loaded EfPiwi molecules, the capture oligo (1.2x nmol of guide RNA used) was bound to High Capacity Neutravidin Resin (Thermo Fisher) in Wash A buffer (30 mM Tris-pH8, 0.1 M KOAc, 2 mM MgOAc, 0.02% CHAPS, 0.5 mM TCEP) for 30 minutes at 4 °C, following which unbound oligo was washed off using Wash A. EfPiwi-guide complex was captured by incubating with Neutravidin resin, containing immobilized capture oligo, at room temperature for 30 minutes with rocking. The resin was then washed three times with 10 ml each Wash A followed by four washes with 10 ml Wash B (30 mM Tris-pH8, 2 M KOAc, 2 mM MgOAc, 0.02% CHAPS, 0.5 mM TCEP) and then three times with 10 ml Wash C (30 mM Tris-pH8, 1 M KOAc, 2 mM MgOAc, 0.04% CHAPS, 0.5 mM TCEP) at 4 °C. Finally, the resin was resuspended in 4-6 ml Wash C, biotinylated elution oligo/competitor DNA was added (2x the amount in nmol of capture oligo used), and the EfPiwi-guide complex was eluted with rocking at room temperature for 2 hours. To remove excess contaminating biotinylated elution oligo, the sample was added to fresh High Capacity Neutravidin resin in Wash C buffer, and the flow through was collected. EfPiwi-guide complex was then dialyzed overnight at 4°C into Q buffer (20 mM Tris pH8, 150 mM NaCl, 0.5 mM TCEP).

EfPiwi-guide complex was incubated with Q Sepharose Fast Flow anion exchange resin (GE Healthcare) at room temperature to remove any excess oligo and the resin was washed with Q wash buffer (20 mM Tris pH8, 250 mM NaCl, 0.5 mM TCEP). Purified EfPiwi-guide complex from the Q flow through was concentrated, had glycerol added to 5-10%, and was flash frozen in liquid nitrogen and stored at -80°C. Concentration of EfPiwi complexes was determined using a Bradford assay with bovine serum albumin (Thermo Fisher) as a standard.

### Grid preparation for cryo-EM

Ternary EfPiwi-piRNA-target complex was formed by adding 1.2 molar equivalents of target RNA to purified EfPiwi-guide complex and incubated on ice for 10 minutes. 3.5 µl EfPiwi-guide complex at 2.5 mg/ml with or without target RNA was added onto freshly plasma cleaned (75% nitrogen, 25% oxygen atmosphere at 15 W for 7 seconds in Solarus plasma cleaner, Gatan) 300 mesh holey gold grids (UltrAuFoil R1.2/1.3, Quantifoil). Excess samples were removed from grids by blotting with Whatman No.1 filter paper for 5-7 s. Samples were immediately vitrified by plunge freezing in liquid-ethane at -179°C using a manual plunge freezing device. Grid vitrification process was performed in a cold room maintained at 4°C with relative humidity between 95-98% to prevent drying up of samples on grids due to evaporation.

### Cryo-EM data acquisition

Cryo-EM data acquisition was performed on a 200kV Talos Arctica (Thermo Fisher Scientific) transmission electron microscope. Micrographs were acquired using a K2 Summit (Gatan) direct electron detector, operated in electron-counting mode, using the automated data collection software Leginon (Suloway et al., 2005) by image shift-based movements from the center of four adjacent holes to target the center of each hole for exposures. Each micrograph for the EfPiwi-piRNA complex was collected as 48 dose-fractionated movie frames over 12 s and with a cumulative electron exposure of 47.33 e^-^/Å^2^. For the EfPiwi-piRNA-target complex, each micrograph was acquired as 64 dose-fractionated movie frames over 16 s with a cumulative electron exposure of 47.33 e^-^/Å^2^. Both data sets were collected at a nominal magnification of 36kx, corresponding to 1.15 Å/pixel on the detector, with random nominal defocus values varying between 1 µm and 1.6 µm. 1,765 and 1,881 micrographs were collected for the EfPiwi-piRNA and EfPIWI-piRNA-target complexes, respectively.

### Image processing and 3D reconstruction

Beam-induced motion correction and radiation damage compensation over spatial frequencies (dose-weighting) of the raw movies, was performed using UCSF MotionCor2 (Zheng et al., 2017) implemented in the Appion (Lander et al., 2009) image processing workflow. Motion corrected, summed micrographs were imported into the RELION 2.0 (Kimanius et al., 2016) data processing pipeline (**Fig S1 B-K**). Contrast Transfer Function (CTF) parameters for these micrographs were estimated using CTFFind4 (Rohou and Grigorieff, 2015). Laplacian of Gaussian based automated particle picking program in RELION was used for picking 3,280,351 and 2,551,046 particles from the EfPiwi-piRNA and EfPiwi-piRNA-target micrographs, respectively. Picked particles were extracted from the micrographs with a 160 pixel box and subjected to 2D classification in RELION. After discarding particles belonging to classes containing non-particle features, aggregates and low-resolution features, new stacks of particles from 2D classes containing different orientations of the complexes and high-resolution features were created. A subset of 2D classes from the EfPiwi-piRNA-target dataset resolved features corresponding to a smaller subcomplex with an extension of an RNA duplex. The 608,488 particles belonging to these classes were isolated into a new particle-stack for further processing. To serve as an initial model for 3D analyses of the EfPiwi-piRNA and EfPiwi-piRNA-target complexes, a 40 Å low pass filtered map was generated from Siwi crystal structure (PDB ID 5GUH) using the molmap function in UCSF Chimera (Peterson et al., 2004, Goddard et al., 2007). An initial model for 3D analyses of the smaller subcomplex particles was generated using the Cryosparc v1 (Brubaker et al., 2017, Punjani et al., 2017) *ab initio* reconstruction program. The selected particle stacks corresponding to the three distinct complexes were subjected to multiple iterations of 3D classification in RELION and particles belonging to the most well-resolved 3D class for each complex were selected for downstream 3D processing. After 3D classification, 125,041, 118,493, and 116,655 particles from the best resolved class for the EfPiwi-piRNA, EfPiwi-piRNA-target, and the smaller subcomplex, respectively, were re-extracted with 160 pixels box from the respective micrographs with re-centered coordinates. These particles were then subjected to 3D refinement in RELION. 3D binary masks for refinement were generated using 15 Å low-pass filtered selected class volume for each of the complexes with a 5-pixel expansion and 8-pixel Gaussian fall-off in RELION. The final reconstructed map for the EfPiwi-piRNA, EfPiwi-piRNA-target and the smaller subcomplex were at 3.8 Å, 3.5 Å and 8.6 Å (at Fourier Shell Correlation value of 0.143), respectively (**Fig. S1C,F,H,K**). Local resolution for these complexes were determined using the local resolution estimation program in RELION (**Fig. S1D,I**) (Table S1) and the local-resolution based filtered maps were used for atomic model building. Maps for EfPiwi-piRNA and EfPiwi-piRNA-target complexes were trimmed to box size of 90 pixels for deposition into EM Data Bank. Directional Fourier Shell Correlations (FSCs) for the reconstructed maps (**Fig. S1F,K**) were estimated using the 3DFSC server (www.3dfsc.salk.edu) (Baldwin et al., 2017)

### Model building and refinement

An initial model for the EfPiwi was obtained by threading the EfPiwi primary sequence onto the Siwi crystal structure (PDB ID 5GUH) using SWISS-MODEL (Waterhouse, Bertoni et al. 2018). Discrete domains from this model were then docked into the EfPiwi-piRNA reconstruction using UCSF Chimera (Peterson et al., 2004, Goddard et al., 2007), followed by manual docking and model building using Coot (Emsley et al., 2010). The EfPiwi-guide-target model was built in a similar fashion, using the EfPiwi-guide structure as an initial model. The models were refined through iterative rounds of manual building and fixing of geometric and rotameric outliers in Coot (Emsley et al., 2010) and real-space refinement optimizing global minimization, atomic displacement parameters, and local grid search using PHENIX (Afonine et al., 2012). In base pairing between guide and target RNAs was maintained by including hydrogen atoms in the EfPiwi-guide-target model. Most residues within the N domain of both models were truncated to alanine at the end of refinement to reflect a lack of supporting cryo-EM density. Model validation were performed using Molprobity (*molprobity.biochem.duke.edu*) (Chen et al., 2010) and PDB validation servers (www.wwpdb.org) (Berman et al., 2007) (**Table S1**). Structural figures were made using PyMOL (Schrödinger, LLC) or UCSF Chimera (Peterson et al., 2004, Goddard et al., 2007).

### Equilibrium binding assays

Equilibrium dissociation constants were determined as described previously (Schirle, Sheu-Gruttadauria et al. 2014). Loaded EfPiwi or hAgo2 samples were incubated with ^32^P 5′-radiolabeled target RNA in binding reaction buffer (30 mM Tris pH 8.0, 100 mM potassium acetate, 0.5 mM TCEP, 0.005% (v/v) NP-40, 0.01 mg/mL baker’s yeast tRNA), in a reaction volume of 100 μl at room temperature. EfPiwi-guide or hAgo2-guide complex concentrations ranged from 1 pM to 25 nM. Loaded EfPiwi or hAgo2 complexes were incubated with 0.1 nM radiolabeled targets for 45 min.

Using a dot-blot apparatus (GE Healthcare), protein-RNA complexes were immobilized on Protran nitrocellulose membrane (0.45 μm pore size, Whatman, GE Healthcare) and unbound RNA immobilized on Hybond Nylon membrane (Amersham, GE Healthcare). Membranes were stacked so that sample is first pulled through the protein-binding membrane, and any excess unbound RNA passes through and binds to the RNA-binding membrane. Samples were applied with vacuum and then washed using 100 μl of ice-cold wash buffer (30 mM Tris pH 8.0, 0.1 M potassium acetate, 0.5 mM TCEP). Membranes were air-dried and visualized by phosphorimaging. Quantification of signal was performed using ImageQuant TL (GE Healthcare).

### Target dissociation assays

Target dissociation rates were determined by incubating guide-loaded EfPiwi and hAgo2 samples with 0.1 nM ^32^P 5′-radiolabeled target RNA in binding reaction buffer (30 mM Tris pH 8.0, 100 mM potassium acetate, 0.5 mM TCEP, 0.005% (v/v) NP-40, 0.01 mg/mL baker’s yeast tRNA) in a single large reaction with a volume of 100 μl per time point planned for the experiment (e.g. 1000 ul for 10 time points) at room temperature for 60 minutes. The concentration of hAgo2 and EfPiwi complexes was 5 nM.

After equilibration, a zero-time point was taken by applying 100 ul of the reaction to the dot-blot apparatus under vacuum and was followed with 100 μl of ice-cold wash buffer (30 mM Tris pH 8.0, 0.1 M potassium acetate, 0.5 mM TCEP). The reaction was started by addition of unlabeled target RNA in large excess, to a final concentration of 300 nM. Aliquots of 100 μl were taken at indicated time points and applied to a dot-blot apparatus under vacuum and followed with 100 μl of ice-cold wash buffer. Time points ranged from 0.25-100 minutes. Membranes were air-dried and visualized by phosphorimaging. Quantification of signal was performed using ImageQuant TL (GE Healthcare).

### Target association assays

Target association rates were determined by incubating 5 pM ^32^P 5′-radiolabeled target RNA in binding reaction buffer (30 mM Tris pH 8.0, 100 mM potassium acetate, 0.5 mM TCEP, 0.005% (v/v) NP-40, 0.01 mg/mL baker’s yeast tRNA) in a single large reaction with a volume of 100 μl per time point planned for the experiment (e.g. 1000 ul for 10 time points) at room temperature for 15 minutes.

A zero-time point was taken by applying 100 μl of the reaction to the dot-blot apparatus under vacuum and was followed with 100 μl of ice-cold wash buffer (30 mM Tris pH 8.0, 0.1 M potassium acetate, 0.5 mM TCEP). The association reaction was started by addition of guide-loaded EfPiwi or hAgo2 at concentrations ranging from 0.05-15 nM. Aliquots of 100 μl were taken at indicated time points and applied to a dot-blot apparatus under vacuum and followed with 100 μl of ice-cold wash buffer as before. Time points ranged from 0.25-15 minutes. Membranes were air-dried and visualized by phosphorimaging. Quantification of signal was performed using ImageQuant TL (GE Healthcare).

### EfPiwi and hAgo2 slicing assays

Purified EfPiwi-guide or hAgo2-guide complexes (∼100 nM, final concentration) were incubated at 37 °C with complementary 5′-^32^P-labeled target RNAs (∼10 nM, final concentrations) in reaction buffer composed of 30 mM Tris pH 8.0, 2 mM MgCl_2_, 0.5 mM TCEP, and 0.01 mg/mL baker’s yeast tRNA. Target cleavage was stopped at various times by mixing aliquots of each reaction with an equal volume of denaturing gel loading buffer (98% w/v formamide, 0.025% xylene cyanol, 0.025% w/v bromophenol blue, 10 mM EDTA pH 8.0). Intact and cleaved target RNAs were resolved by denaturing PAGE (15%) and visualized by phosphorimaging. Quantification of signal was performed using ImageQuant TL (GE Healthcare).

Adjustments were made to this protocol to determine which divalent cations were catalytic with EfPiwi, by replacing 2 mM MgCl_2_ with 2 mM MnCl_2_, CoCl_2_, CaCl_2_, or NiCl_2_. Similar adjustments were made to titrate in MgCl_2_ or MnCl_2_.

To determine the target length requirement for slicing, 100 nM protein-guide complex and 1 nM radiolabeled target RNA of complementarity g2-g15, g2-g16, g2-g17, g2-g18, g2-g19, g2-g20, or g2-g21 were combined in buffer with 2 mM each MgCl_2_ and MnCl_2_. Reactions proceeded for 1 hour at 37 °C before having an equal volume of denaturing gel loading buffer added to stop. These reactions were run on denaturing PAGE (15%) and analyzed as described above.

To determine the effects of the triplet mismatches on slicing, 100 nM protein-guide complex and 1 nM radiolabeled target RNA with mismatches from g2-g4, g5-g7, g8-g10, g11-g13, g14-g16, g17-g19, or g20-g22 were combined in buffer with 2 mM each MgCl_2_ and MnCl_2_. Reactions proceeded for 1 hour at 37 °C before having an equal volume of denaturing gel loading buffer added to stop. These reactions were run on denaturing PAGE (15%) and analyzed as described above.

## QUANTIFICATION AND STATISTICAL ANALYSIS

### Target RNA equilibrium binding experiments

The fraction of target RNA bound was calculated as the ratio of bound to total (bound + free) target RNA for various concentrations of Ago2**–**guide or EfPiwi-guide complexes. RNA was quantified by phosphorimaging using ImageQuant TL (GE Healthcare). N=3 for each binding experiment, where N represents the number of experimental replicates. Dissociation constants calculated using Prism version 8.0 (GraphPad Software, Inc.), using the One-Site Specific Binding equation. The equation is as follows:

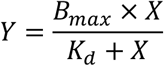

Where Y is specific binding, X is the concentration of the radioligand, B_max_ is the maximum binding, and K_d_ is the dissociation constant. Data between targets were normalized by dividing fraction bound by experimentally determined B_max_ values for individual target RNAs.

### Target association and dissociation assays

The fraction of target RNA bound was calculated as the ratio of bound to total (bound + free) target RNA for various concentrations of Ago2-guide or EfPiwi-guide complexes. RNA was quantified by phosphorimaging using ImageQuant TL (GE Healthcare). N=3 for each binding experiment, where N represents number of experimental replications. Dissociation rates were calculated by fitting data to a one-phase exponential decay curve using Prism version 8.0 (GraphPad Software, Inc.). The equation is as follows:

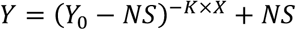

Where X is time, Y is the fraction of the target RNA bound, Y_0_ is Y at time 0, NS is bound fraction at very long times, and K is the rate constant.

Because dissociation was very slow compared to association, and the target RNA concentrations were always at least 10-fold less than total EfPiwi-piRNA concentration, association was treated as an irreversible pseudo first-order system. Data were fit to a one-phase exponential curve to determine an observed binding rate constant (*k*_*obs*_). Because the *k*_*off*_ is constant regardless of the protein concentration (units of min^-1^), normalization between samples of varying concentration was not required. However, the observed *k*_*on*_ changed as a function of protein concentration, and therefore *k*_*on*_ was calculated by dividing *k*_*obs*_ by the concentration of the EfPiwi-piRNA complex used in the experiment.

### Target cleavage assays

The fraction of cleaved target RNA was determined by calculating the ratio of cleaved to total (cleaved + uncleaved) target RNA. For time courses, rate of cleavage was calculated by fitting data to a one-phase exponential decay curve using Prism version 8.0 (GraphPad Software, Inc.). Equation is as above.

## References

Alie, A., T. Hayashi, I. Sugimura, M. Manuel, W. Sugano, A. Mano, N. Satoh, K. Agata and N. Funayama (2015). “The ancestral gene repertoire of animal stem cells.” Proc Natl Acad Sci U S A 112(51): E7093–7100.

Aravin, A., D. Gaidatzis, S. Pfeffer, M. Lagos-Quintana, P. Landgraf, N. Iovino, P. Morris, M. J. Brownstein, S. Kuramochi-Miyagawa, T. Nakano, M. Chien, J. J. Russo, J. Ju, R. Sheridan, C. Sander, M. Zavolan and T. Tuschl (2006). “A novel class of small RNAs bind to MILI protein in mouse testes.” Nature 442(7099): 203–207.

Aravin, A. A., R. Sachidanandam, D. Bourc’his, C. Schaefer, D. Pezic, K. F. Toth, T. Bestor and G. J. Hannon (2008). “A piRNA pathway primed by individual transposons is linked to de novo DNA methylation in mice.” Mol Cell 31(6): 785–799.

Aravin, A. A., R. Sachidanandam, A. Girard, K. Fejes-Toth and G. J. Hannon (2007). “Developmentally regulated piRNA clusters implicate MILI in transposon control.” Science 316(5825): 744–747.

Bartel, D. P. (2018). “Metazoan MicroRNAs.” Cell 173(1): 20–51.

Brennecke, J., A. A. Aravin, A. Stark, M. Dus, M. Kellis, R. Sachidanandam and G. J. Hannon (2007). “Discrete small RNA-generating loci as master regulators of transposon activity in Drosophila.” Cell 128(6): 1089–1103.

Brennecke, J., C. D. Malone, A. A. Aravin, R. Sachidanandam, A. Stark and G. J. Hannon (2008). “An epigenetic role for maternally inherited piRNAs in transposon silencing.” Science 322(5906): 1387–1392.

Brennecke, J., A. Stark, R. B. Russell and S. M. Cohen (2005). “Principles of microRNA-target recognition.” PLoS Biol 3(3): e85.

Carmell, M. A., Z. Xuan, M. Q. Zhang and G. J. Hannon (2002). “The Argonaute family: tentacles that reach into RNAi, developmental control, stem cell maintenance, and tumorigenesis.” Genes Dev 16(21): 2733–2742.

Chandradoss, S. D., N. T. Schirle, M. Szczepaniak, I. J. MacRae and C. Joo (2015). “A Dynamic Search Process Underlies MicroRNA Targeting.” Cell 162(1): 96–107.

De Fazio, S., N. Bartonicek, M. Di Giacomo, C. Abreu-Goodger, A. Sankar, C. Funaya, C. Antony, P. N. Moreira, A. J. Enright and D. O’Carroll (2011). “The endonuclease activity of Mili fuels piRNA amplification that silences LINE1 elements.” Nature 480(7376): 259–263.

Filipowicz, W. (2005). “RNAi: the nuts and bolts of the RISC machine.” Cell 122(1): 17–20.

Flores-Jasso, C. F., W. E. Salomon and P. D. Zamore (2013). “Rapid and specific purification of Argonaute-small RNA complexes from crude cell lysates.” RNA 19(2): 271–279.

Funayama, N., M. Nakatsukasa, K. Mohri, Y. Masuda and K. Agata (2010). “Piwi expression in archeocytes and choanocytes in demosponges: insights into the stem cell system in demosponges.” Evol Dev 12(3): 275–287.

Gainetdinov, I., C. Colpan, A. Arif, K. Cecchini and P. D. Zamore (2018). “A Single Mechanism of Biogenesis, Initiated and Directed by PIWI Proteins, Explains piRNA Production in Most Animals.” Mol Cell 71(5): 775–790 e775.

Girard, A., R. Sachidanandam, G. J. Hannon and M. A. Carmell (2006). “A germline-specific class of small RNAs binds mammalian Piwi proteins.” Nature 442(7099): 199–202.

Goh, W. S., I. Falciatori, O. H. Tam, R. Burgess, O. Meikar, N. Kotaja, M. Hammell and G. J. Hannon (2015). “piRNA-directed cleavage of meiotic transcripts regulates spermatogenesis.” Genes Dev 29(10): 1032–1044.

Gou, L. T., P. Dai, J. H. Yang, Y. Xue, Y. P. Hu, Y. Zhou, J. Y. Kang, X. Wang, H. Li, M. M. Hua, S. Zhao, S. D. Hu, L. G. Wu, H. J. Shi, Y. Li, X. D. Fu, L. H. Qu, E. D. Wang and M. F. Liu (2014). “Pachytene piRNAs instruct massive mRNA elimination during late spermiogenesis.” Cell Res 24(6): 680–700.

Grimson, A., K. K. Farh, W. K. Johnston, P. Garrett-Engele, L. P. Lim and D. P. Bartel (2007). “MicroRNA targeting specificity in mammals: determinants beyond seed pairing.” Mol Cell 27(1): 91–105.

Grimson, A., M. Srivastava, B. Fahey, B. J. Woodcroft, H. R. Chiang, N. King, B. M. Degnan, D. S. Rokhsar and D. P. Bartel (2008). “Early origins and evolution of microRNAs and Piwi-interacting RNAs in animals.” Nature 455(7217): 1193–1197.

Grivna, S. T., E. Beyret, Z. Wang and H. Lin (2006). “A novel class of small RNAs in mouse spermatogenic cells.” Genes Dev 20(13): 1709–1714.

Grosswendt, S., A. Filipchyk, M. Manzano, F. Klironomos, M. Schilling, M. Herzog, E. Gottwein and N. Rajewskys (2014). “Unambiguous Identification of miRNA:Target Site Interactions by Different Types of Ligation Reactions.” Mol Cell 54(6): 1042–1054.

Grosswendt, S., A. Filipchyk, M. Manzano, F. Klironomos, M. Schilling, M. Herzog, E. Gottwein and N. Rajewsky (2014). “Unambiguous identification of miRNA:target site interactions by different types of ligation reactions.” Mol Cell 54(6): 1042–1054.

Gunawardane, L. S., K. Saito, K. M. Nishida, K. Miyoshi, Y. Kawamura, T. Nagami, H. Siomi and M. C. Siomi (2007). “A slicer-mediated mechanism for repeat-associated siRNA 5’ end formation in Drosophila.” Science 315(5818): 1587–1590.

Halbach, R., P. Miesen, J. Joosten, E. Taskopru, I. Rondeel, B. Pennings, C. B. F. Vogels, S. H. Merkling, C. J. Koenraadt, L. Lambrechts and R. P. van Rij (2020). “A satellite repeat-derived piRNA controls embryonic development of Aedes.” Nature 580(7802): 274–277.

Haley, B. and P. D. Zamore (2004). “Kinetic analysis of the RNAi enzyme complex.” Nat Struct Mol Biol 11(7): 599–606.

Han, B. W., W. Wang, C. Li, Z. Weng and P. D. Zamore (2015). “piRNA-guided transposon cleavage initiates Zucchini-dependent, phased piRNA production.” Science 348(6236): 817–821.

Herzog, V. A., B. Reichholf, T. Neumann, P. Rescheneder, P. Bhat, T. R. Burkard, W. Wlotzka, A. von Haeseler, J. Zuber and S. L. Ameres (2017). “Thiol-linked alkylation of RNA to assess expression dynamics.” Nat Methods 14(12): 1198–1204.

Homolka, D., R. R. Pandey, C. Goriaux, E. Brasset, C. Vaury, R. Sachidanandam, M. O. Fauvarque and R. S. Pillai (2015). “PIWI Slicing and RNA Elements in Precursors Instruct Directional Primary piRNA Biogenesis.” Cell Rep 12(3): 418–428.

Houwing, S., L. M. Kamminga, E. Berezikov, D. Cronembold, A. Girard, H. van den Elst, D. V. Filippov, H. Blaser, E. Raz, C. B. Moens, R. H. Plasterk, G. J. Hannon, B. W. Draper and R. F. Ketting (2007). “A role for Piwi and piRNAs in germ cell maintenance and transposon silencing in Zebrafish.” Cell 129(1): 69–82.

Jehn, J., D. Gebert, F. Pipilescu, S. Stern, J. S. T. Kiefer, C. Hewel and D. Rosenkranz (2018). “PIWI genes and piRNAs are ubiquitously expressed in mollusks and show patterns of lineage-specific adaptation.” Commun Biol 1: 137.

Jones, D. T. and D. Cozzetto (2015). “DISOPRED3: precise disordered region predictions with annotated protein-binding activity.” Bioinformatics 31(6): 857–863.

Juliano, C. E., A. Reich, N. Liu, J. Gotzfried, M. Zhong, S. Uman, R. A. Reenan, G. M. Wessel, R. E. Steele and H. Lin (2014). “PIWI proteins and PIWI-interacting RNAs function in Hydra somatic stem cells.” Proc Natl Acad Sci U S A 111(1): 337–342.

Kawaoka, S., N. Hayashi, Y. Suzuki, H. Abe, S. Sugano, Y. Tomari, T. Shimada and S. Katsuma (2009). “The Bombyx ovary-derived cell line endogenously expresses PIWI/PIWI-interacting RNA complexes.” RNA 15(7): 1258–1264.

Kaya, E., K. W. Doxzen, K. R. Knoll, R. C. Wilson, S. C. Strutt, P. J. Kranzusch and J. A. Doudna (2016). “A bacterial Argonaute with noncanonical guide RNA specificity.” Proc Natl Acad Sci U S A 113(15): 4057–4062.

Khurana, J. S., J. Wang, J. Xu, B. S. Koppetsch, T. C. Thomson, A. Nowosielska, C. Li, P. D. Zamore, Z. Weng and W. E. Theurkauf (2011). “Adaptation to P element transposon invasion in Drosophila melanogaster.” Cell 147(7): 1551–1563.

Kiuchi, T., H. Koga, M. Kawamoto, K. Shoji, H. Sakai, Y. Arai, G. Ishihara, S. Kawaoka, S. Sugano, T. Shimada, Y. Suzuki, M. G. Suzuki and S. Katsuma (2014). “A single female-specific piRNA is the primary determiner of sex in the silkworm.” Nature 509(7502): 633–636.

Klum, S. M., S. D. Chandradoss, N. T. Schirle, C. Joo and I. J. MacRae (2018). “Helix-7 in Argonaute2 shapes the microRNA seed region for rapid target recognition.” EMBO J 37(1): 75–88.

Krek, A., D. Grun, M. N. Poy, R. Wolf, L. Rosenberg, E. J. Epstein, P. MacMenamin, I. da Piedade, K. C. Gunsalus, M. Stoffel and N. Rajewsky (2005). “Combinatorial microRNA target predictions.” Nat Genet 37(5): 495–500.

Kuramochi-Miyagawa, S., T. Watanabe, K. Gotoh, Y. Totoki, A. Toyoda, M. Ikawa, N. Asada, K. Kojima, Y. Yamaguchi, T. W. Ijiri, K. Hata, E. Li, Y. Matsuda, T. Kimura, M. Okabe, Y. Sakaki, H. Sasaki and T. Nakano (2008). “DNA methylation of retrotransposon genes is regulated by Piwi family members MILI and MIWI2 in murine fetal testes.” Genes Dev 22(7): 908–917.

Lai, E. C. (2002). “Micro RNAs are complementary to 3’ UTR sequence motifs that mediate negative post-transcriptional regulation.” Nat Genet 30(4): 363–364.

Lai, E. C. and J. W. Posakony (1998). “Regulation of Drosophila neurogenesis by RNA:RNA duplexes?” Cell 93(7): 1103–1104.

Lau, N. C., A. G. Seto, J. Kim, S. Kuramochi-Miyagawa, T. Nakano, D. P. Bartel and R. E. Kingston (2006). “Characterization of the piRNA complex from rat testes.” Science 313(5785): 363–367.

Le Thomas, A., A. K. Rogers, A. Webster, G. K. Marinov, S. E. Liao, E. M. Perkins, J. K. Hur, A. A. Aravin and K. F. Toth (2013). “Piwi induces piRNA-guided transcriptional silencing and establishment of a repressive chromatin state.” Genes Dev 27(4): 390–399.

Lewis, B. P., I. H. Shih, M. W. Jones-Rhoades, D. P. Bartel and C. B. Burge (2003). “Prediction of mammalian microRNA targets.” Cell 115(7): 787–798.

Lim, L. P., N. C. Lau, P. Garrett-Engele, A. Grimson, J. M. Schelter, J. Castle, D. P. Bartel, P. S. Linsley and J. M. Johnson (2005). “Microarray analysis shows that some microRNAs downregulate large numbers of target mRNAs.” Nature 433(7027): 769–773.

Lin, H. and A. C. Spradling (1997). “A novel group of pumilio mutations affects the asymmetric division of germline stem cells in the Drosophila ovary.” Development 124(12): 2463–2476.

Matsumoto, N., H. Nishimasu, K. Sakakibara, K. M. Nishida, T. Hirano, R. Ishitani, H. Siomi, M. C. Siomi and O. Nureki (2016). “Crystal Structure of Silkworm PIWI-Clade Argonaute Siwi Bound to piRNA.” Cell 167(2): 484–497 e489.

Miesen, P., J. Joosten and R. P. van Rij (2016). “PIWIs Go Viral: Arbovirus-Derived piRNAs in Vector Mosquitoes.” PLoS Pathog 12(12): e1006017.

Mohn, F., D. Handler and J. Brennecke (2015). “piRNA-guided slicing specifies transcripts for Zucchini-dependent, phased piRNA biogenesis.” Science 348(6236): 812–817.

Nakanishi, K., M. Ascano, T. Gogakos, S. Ishibe-Murakami, A. A. Serganov, D. Briskin, P. Morozov, T. Tuschl and D. J. Patel (2013). “Eukaryote-specific insertion elements control human ARGONAUTE slicer activity.” Cell Rep 3(6): 1893–1900.

Nakanishi, K., D. E. Weinberg, D. P. Bartel and D. J. Patel (2012). “Structure of yeast Argonaute with guide RNA.” Nature 486(7403): 368–374.

Nishida, K. M., Y. W. Iwasaki, Y. Murota, A. Nagao, T. Mannen, Y. Kato, H. Siomi and M. C. Siomi (2015). “Respective functions of two distinct Siwi complexes assembled during PIWI-interacting RNA biogenesis in Bombyx germ cells.” Cell Rep 10(2): 193–203.

Nott, T. J., E. Petsalaki, P. Farber, D. Jervis, E. Fussner, A. Plochowietz, T. D. Craggs, D. P. Bazett-Jones, T. Pawson, J. D. Forman-Kay and A. J. Baldwin (2015). “Phase transition of a disordered nuage protein generates environmentally responsive membraneless organelles.” Mol Cell 57(5): 936–947.

Nozawa, M., M. Fujimi, C. Iwamoto, K. Onizuka, N. Fukuda, K. Ikeo and T. Gojobori (2016). “Evolutionary Transitions of MicroRNA-Target Pairs.” Genome Biol Evol 8(5): 1621–1633.

Ozata, D. M., I. Gainetdinov, A. Zoch, D. O’Carroll and P. D. Zamore (2019). “PIWI-interacting RNAs: small RNAs with big functions.” Nat Rev Genet 20(2): 89–108.

Park, M. S., R. Araya-Secchi, J. A. Brackbill, H. D. Phan, A. C. Kehling, E. W. Abd El-Wahab, D. M. Dayeh, M. Sotomayor and K. Nakanishi (2019). “Multidomain Convergence of Argonaute during RISC Assembly Correlates with the Formation of Internal Water Clusters.” Mol Cell 75(4): 725–740 e726.

Park, M. S., H. D. Phan, F. Busch, S. H. Hinckley, J. A. Brackbill, V. H. Wysocki and K. Nakanishi (2017). “Human Argonaute3 has slicer activity.” Nucleic Acids Res 45(20): 11867–11877.

Parker, J. S., E. A. Parizotto, M. Wang, S. M. Roe and D. Barford (2009). “Enhancement of the seed-target recognition step in RNA silencing by a PIWI/MID domain protein.” Mol Cell 33(2): 204–214.

Parker, J. S., S. M. Roe and D. Barford (2005). “Structural insights into mRNA recognition from a PIWI domain-siRNA guide complex.” Nature 434(7033): 663–666.

Rajasethupathy, P., I. Antonov, R. Sheridan, S. Frey, C. Sander, T. Tuschl and E. R. Kandel (2012). “A role for neuronal piRNAs in the epigenetic control of memory-related synaptic plasticity.” Cell 149(3): 693–707.

Reddien, P. W., N. J. Oviedo, J. R. Jennings, J. C. Jenkin and A. Sanchez Alvarado (2005). “SMEDWI-2 is a PIWI-like protein that regulates planarian stem cells.” Science 310(5752): 1327–1330.

Reuter, M., P. Berninger, S. Chuma, H. Shah, M. Hosokawa, C. Funaya, C. Antony, R. Sachidanandam and R. S. Pillai (2011). “Miwi catalysis is required for piRNA amplification-independent LINE1 transposon silencing.” Nature 480(7376): 264–267.

Rinkevich, Y., A. Rosner, C. Rabinowitz, Z. Lapidot, E. Moiseeva and B. Rinkevich (2010). “Piwi positive cells that line the vasculature epithelium, underlie whole body regeneration in a basal chordate.” Dev Biol 345(1): 94–104.

Rivas, F. V., N. H. Tolia, J. J. Song, J. P. Aragon, J. Liu, G. J. Hannon and L. Joshua-Tor (2005). “Purified Argonaute2 and an siRNA form recombinant human RISC.” Nat Struct Mol Biol 12(4): 340–349.

Rouget, C., C. Papin, A. Boureux, A. C. Meunier, B. Franco, N. Robine, E. C. Lai, A. Pelisson and M. Simonelig (2010). “Maternal mRNA deadenylation and decay by the piRNA pathway in the early Drosophila embryo.” Nature 467(7319): 1128–1132.

Rubin, G. M., M. G. Kidwell and P. M. Bingham (1982). “The molecular basis of P-M hybrid dysgenesis: the nature of induced mutations.” Cell 29(3): 987–994.

Salomon, W. E., S. M. Jolly, M. J. Moore, P. D. Zamore and V. Serebrov (2015). “Single-Molecule Imaging Reveals that Argonaute Reshapes the Binding Properties of Its Nucleic Acid Guides.” Cell 162(1): 84–95.

Schirle, N. T. and I. J. MacRae (2012). “The crystal structure of human Argonaute2.” Science 336(6084): 1037–1040.

Schirle, N. T., J. Sheu-Gruttadauria and I. J. MacRae (2014). “Structural basis for microRNA targeting.” Science 346(6209): 608–613.

Schwarz, D. S., Y. Tomari and P. D. Zamore (2004). “The RNA-induced silencing complex is a Mg2+-dependent endonuclease.” Curr Biol 14(9): 787–791.

Shen, E. Z., H. Chen, A. R. Ozturk, S. Tu, M. Shirayama, W. Tang, Y. H. Ding, S. Y. Dai, Z. Weng and C. C. Mello (2018). “Identification of piRNA Binding Sites Reveals the Argonaute Regulatory Landscape of the C. elegans Germline.” Cell 172(5): 937–951 e918.

Sheng, G., H. Zhao, J. Wang, Y. Rao, W. Tian, D. C. Swarts, J. van der Oost, D. J. Patel and Y. Wang (2014). “Structure-based cleavage mechanism of Thermus thermophilus Argonaute DNA guide strand-mediated DNA target cleavage.” Proc Natl Acad Sci U S A 111(2): 652–657.

Sheu-Gruttadauria, J., P. Pawlica, S. M. Klum, S. Wang, T. A. Yario, N. T. Schirle Oakdale, J. A. Steitz and I. J. MacRae (2019). “Structural Basis for Target-Directed MicroRNA Degradation.” Mol Cell.

Sheu-Gruttadauria, J., P. Pawlica, S. M. Klum, S. Wang, T. A. Yario, N. T. Schirle Oakdale, J. A. Steitz and I. J. MacRae (2019). “Structural Basis for Target-Directed MicroRNA Degradation.” Mol Cell 75(6): 1243–1255 e1247.

Sheu-Gruttadauria, J., Y. Xiao, L. F. Gebert and I. J. MacRae (2019). “Beyond the seed: structural basis for supplementary microRNA targeting by human Argonaute2.” EMBO J 38(13): e101153.

Sheu-Gruttadauria, J., Y. Xiao, L. F. Gebert and I. J. MacRae (2019). “Beyond the seed: structural basis for supplementary microRNA targeting by human Argonaute2.” EMBO J.

Sienski, G., D. Donertas and J. Brennecke (2012). “Transcriptional silencing of transposons by Piwi and maelstrom and its impact on chromatin state and gene expression.” Cell 151(5): 964–980.

Simkin, A., R. Geissler, A. B. R. McIntyre and A. Grimson (2020). “Evolutionary dynamics of microRNA target sites across vertebrate evolution.” PLoS Genet 16(2): e1008285.

Swarts, D. C., J. W. Hegge, I. Hinojo, M. Shiimori, M. A. Ellis, J. Dumrongkulraksa, R. M. Terns, M. P. Terns and J. van der Oost (2015). “Argonaute of the archaeon Pyrococcus furiosus is a DNA-guided nuclease that targets cognate DNA.” Nucleic Acids Res 43(10): 5120–5129.

Tian, Y., D. K. Simanshu, J. B. Ma and D. J. Patel (2011). “Structural basis for piRNA 2’-O-methylated 3’-end recognition by Piwi PAZ (Piwi/Argonaute/Zwille) domains.” Proc Natl Acad Sci U S A 108(3): 903–910.

Tomari, Y. and P. D. Zamore (2005). “Perspective: machines for RNAi.” Genes Dev 19(5): 517–529.

Vagin, V. V., A. Sigova, C. Li, H. Seitz, V. Gvozdev and P. D. Zamore (2006). “A distinct small RNA pathway silences selfish genetic elements in the germline.” Science 313(5785): 320–324.

Wang, J., P. Zhang, Y. Lu, Y. Li, Y. Zheng, Y. Kan, R. Chen and S. He (2019). “piRBase: a comprehensive database of piRNA sequences.” Nucleic Acids Res 47(D1): D175–D180.

Wang, Y., S. Juranek, H. Li, G. Sheng, G. S. Wardle, T. Tuschl and D. J. Patel (2009). “Nucleation, propagation and cleavage of target RNAs in Ago silencing complexes.” Nature 461(7265): 754–761.

Wang, Y., G. Sheng, S. Juranek, T. Tuschl and D. J. Patel (2008). “Structure of the guide-strand-containing argonaute silencing complex.” Nature 456(7219): 209–213.

Waterhouse, A., M. Bertoni, S. Bienert, G. Studer, G. Tauriello, R. Gumienny, F. T. Heer, T. A. P. de Beer, C. Rempfer, L. Bordoli, R. Lepore and T. Schwede (2018). “SWISS-MODEL: homology modelling of protein structures and complexes.” Nucleic Acids Res 46(W1): W296–W303.

Wee, L. M., C. F. Flores-Jasso, W. E. Salomon and P. D. Zamore (2012). “Argonaute divides its RNA guide into domains with distinct functions and RNA-binding properties.” Cell 151(5): 1055–1067.

Wu, P. H., Y. Fu, K. Cecchini, D. M. Ozata, A. Arif, T. Yu, C. Colpan, I. Gainetdinov, Z. Weng and P. D. Zamore (2020). “The evolutionarily conserved piRNA-producing locus pi6 is required for male mouse fertility.” Nat Genet 52(7): 728–739.

Wynant, N., D. Santos and J. Vanden Broeck (2017). “The evolution of animal Argonautes: evidence for the absence of antiviral AGO Argonautes in vertebrates.” Sci Rep 7(1): 9230.

Xiol, J., P. Spinelli, M. A. Laussmann, D. Homolka, Z. Yang, E. Cora, Y. Coute, S. Conn, J. Kadlec, R. Sachidanandam, M. Kaksonen, S. Cusack, A. Ephrussi and R. S. Pillai (2014). “RNA clamping by Vasa assembles a piRNA amplifier complex on transposon transcripts.” Cell 157(7): 1698–1711.

Yamaguchi, S., A. Oe, K. M. Nishida, K. Yamashita, A. Kajiya, S. Hirano, N. Matsumoto, N. Dohmae, R. Ishitani, K. Saito, H. Siomi, H. Nishimasu, M. C. Siomi and O. Nureki (2020). “Crystal structure of Drosophila Piwi.” Nat Commun 11(1): 858.

Yuan, Y. R., Y. Pei, J. B. Ma, V. Kuryavyi, M. Zhadina, G. Meister, H. Y. Chen, Z. Dauter, T. Tuschl and D. J. Patel (2005). “Crystal structure of A. aeolicus argonaute, a site-specific DNA-guided endoribonuclease, provides insights into RISC-mediated mRNA cleavage.” Mol Cell 19(3): 405–419.

Zhang, D., S. Tu, M. Stubna, W. S. Wu, W. C. Huang, Z. Weng and H. C. Lee (2018). “The piRNA targeting rules and the resistance to piRNA silencing in endogenous genes.” Science 359(6375): 587–592.

Zhang, P., J. Y. Kang, L. T. Gou, J. Wang, Y. Xue, G. Skogerboe, P. Dai, D. W. Huang, R. Chen, X. D. Fu, M. F. Liu and S. He (2015). “MIWI and piRNA-mediated cleavage of messenger RNAs in mouse testes.” Cell Res 25(2): 193–207.

Zhu, W., G. M. Pao, A. Satoh, G. Cummings, J. R. Monaghan, T. T. Harkins, S. V. Bryant, S. Randal Voss, D. M. Gardiner and T. Hunter (2012). “Activation of germline-specific genes is required for limb regeneration in the Mexican axolotl.” Dev Biol 370(1): 42–51.

